# Somatosensory-evoked response induces extensive diffusivity and kurtosis changes associated with neural activity in rodents

**DOI:** 10.1101/2024.04.26.591285

**Authors:** Andreea Hertanu, Tommaso Pavan, Ileana O. Jelescu

**Author notes:** Andreea Hertanu, Department of Radiology, Lausanne University Hospital (CHUV) Rue du Bugnon 46, 1011 Lausanne.

## Abstract

Neural tissue microstructure is dynamic during brain activity, presenting changes in cellular morphology and membrane permeability. The sensitivity of diffusion MRI (dMRI) to restrictions and hindrances in the form of cell membranes or subcellular structures enables the exploration of brain activity under a new paradigm, offering a more direct functional contrast than its BOLD counterpart. The current work aims at probing Mean Diffusivity (MD) and Mean Kurtosis (MK) changes and their time-dependence signature across various regions in the rat brain during somatosensory processing and integration, upon unilateral forepaw stimulation. We report a *decrease* in MD in the contralateral primary somatosensory cortex, forelimb region (S1FL), previously ascribed to cellular swelling and increased tortuosity in the extracellular space, paralleled by a positive BOLD response. For the first time, we also report a paired *decrease* in MK during stimulation in S1FL, suggesting increased membrane permeability. This observation was further supported by the reduction in exchange time estimated from the kurtosis time-dependence analyses. Conversely, the secondary somatosensory cortex and subcortical areas, formerly reported as responsive to sensory stimulation in rodents (thalamus, striatum, hippocampal subfields), displayed a marked MD and MK *increase,* paralleled by a weak-to-absent BOLD response. Overall, MD and MK uncovered functional-induced changes with higher sensitivity than BOLD. Although the exact origin of the MD and MK increase is yet to be unraveled, the potential of dMRI to provide complementary functional insights, even below the BOLD detection threshold, has been showcased.

## 1. INTRODUCTION

Neural activity involves the coordinated interplay of neurovascular and neurometabolic coupling mechanisms, where electrical signals in neurons trigger changes in blood flow and metabolism (Sokoloff, 1978). This complex process encompasses action potentials, synaptic transmission, and the integration of excitatory and inhibitory signals, contributing to essential brain functions.

By measuring the regional disparity in oxygen consumption and supply, the blood-oxygenation-level-dependent (BOLD) technique harnesses the interaction between hemodynamic and metabolic factors and provides a surrogate for brain activity (Ogawa et al., 1990). Since its inception, functional Magnetic Resonance Imaging (fMRI) based on the BOLD effect (BOLD-fMRI) has become widely used and applied in the field of neuroscience. Over the years, BOLD-fMRI has been instrumental in investigating various aspects of brain function, from the intricacies of memory and attention to the examination of neurological disorders (Matthews & Jezzard, 2004). Its versatility extends beyond clinical realms, contributing to the exploration of responses to sensory stimuli, decision-making, and social cognition (Morita et al., 2016).

A less well-known coupling linking neural activity and the mechanical tissue properties was illustrated by several groups reporting transient activity-dependent neuronal and glial swelling (Tasaki & Byrne, 1992; Saly & Andrew, 1993; Andrew & Macvicar, 1994; Sun & Wu, 2001; Fraser & Huang, 2004; Chéreau et al., 2017; Costa et al., 2018; Ling et al., 2020; Kwon et al., 2023). The neuromorphological coupling was hypothesized as an underlying mechanism for another type of fMRI contrast based on diffusion MRI (dMRI), thought to provide a more direct mapping of neural activity as compared to BOLD (Aso et al., 2013; Le Bihan et al., 2006). Indeed, the spatial specificity and sensitivity of BOLD-fMRI is a challenging and stimulating debate (Norris, 2012) heightened by nonlinear, time-varying, and partially unknown crosstalk between hemodynamic, metabolic, and neuronal activity (Buxton et al., 1998; Friston et al., 2000; Sundqvist et al., 2022). On the other hand, diffusion functional MRI (dfMRI) (Abe, Tsurugizawa, et al., 2017; Aso et al., 2013; Darquié et al., 2001; Le Bihan et al., 2006; Tsurugizawa et al., 2013) leverages the sensitivity of the diffusion-weighted signal to microstructural features and provides a distinctive approach to probe cellular morphology fluctuations accompanying neural activity.

Owing to methodological hurdles, dfMRI has been subject to controversy over its ability to reflect neuromorphological changes rather than plain vascular modifications (i.e. BOLD contribution) (Bai et al., 2016; Kuroiwa et al., 2014; Miller et al., 2007; Williams et al., 2016). However, recent studies with optimal signal-to-noise ratio (SNR) (Nunes et al., 2019), ultra-high temporal resolution (Nunes et al., 2021) and careful acquisition schemes minimizing vascular confounds (Olszowy et al., 2021) have shown that non-vascular information can be gleaned from dfMRI. Notably, time-courses of apparent diffusion coefficient (ADC) estimated from pairs of diffusion-weighted images acquired in an interleaved fashion, rather than time-courses of diffusion-weighted signals, cancel out most vascular contributions to dfMRI through T_2_-weighting (Darquié et al., 2001; Nunes et al., 2021; Olszowy et al., 2021; Yacoub et al., 2008). The typical reduction in ADC during neuronal activation reported in such dfMRI studies was attributed to increased tortuosity in the extracellular space and cellular swelling and was detected during brain activity even in the absence of neurovascular coupling (Tsurugizawa et al., 2013; Flint et al., 2009; Tirosh & Nevo, 2013; Spees et al., 2013; Lin et al., 2014).

Furthermore, impairment of the water transport mechanisms taking place through the astrocytic aquaporin channels reflected as a decrease in ADC in the living rodent brain (Badaut et al., 2011; Pavlin et al., 2017), while the addition of neural activity inhibitors in rat brain organotypic cortical cultures (Bai et al., 2019) led to a considerable decrease in the trans-membrane active water cycling mechanisms measured through relaxometry.

Time-dependent analyses can provide insights into both tissue microstructure and permeability characteristics (Lee et al., 2020). For instance, at short to intermediate diffusion times, water molecules may not fully explore their environment, leading to non-Gaussian diffusion. In the long-time limit, diffusion becomes Gaussian as the water molecules have interacted with all of the environment equally (so-called coarse-graining). The way mean diffusivity (MD) and mean kurtosis (MK) vary with diffusion time (i.e. the functional form of this time-dependence) informs on the type of structural disorder the molecules are encountering. For example, the functional form associated with 1D-disorder (Novikov et al., 2014) applies to intra-cellular water in axons, dendrites or other cell processes and can provide information on beading or spines. The functional form associated with 2D-3D structural disorder (Novikov et al., 2014) applies to extra-cellular water and can provide information on outer axon diameter or other characteristic lengths of the extracellular space. For a given family of structural disorder, the same functional form will govern both MD and MK time-dependence. Additionally, MK time-dependence may stem from inter-compartmental water exchange, which can be estimated using the Kärger model (Kärger, 1985).

In this context, the goal of our work was to investigate how brain activity changes MD and MK values (at each diffusion time) but potentially also changes their diffusion time-dependence signature, which could provide additional insight into activity-driven microstructure changes. The fast kurtosis method (Hansen, Lund, et al., 2016; Hansen, Shemesh, et al., 2016) facilitated the estimation of MD and MK for multiple diffusion times from a reduced number of diffusion-weighted images, compatible with the brief rest and stimulus intervals typical of task fMRI block designs. The rat forepaw unilateral electrical stimulation was chosen due to its widespread adoption and extensive use as an experimental model for the exploration of brain function along the somatosensory pathways in rodents. We aimed at probing brain-wide MD and MK dynamics, including various cortical and subcortical structures reported in the literature as part of somatosensory processing and integration.

## 2. MATERIALS AND METHODS

### 2.1 Animal Preparation and Anesthesia

Experiments were approved by the local Service for Veterinary Affairs of the canton of Vaud. Ten female Wistar rats (Charles River, France) (236 ± 14g) were scanned on a 14.1T MRI system (Bruker BioSpin), using a home-built surface quadrature transceiver RF-coil. The anesthesia, stimulation and animal preparation protocols were similar to those outlined in a previous study (Reynaud et al., 2019). Isoflurane anesthesia (4% for induction and 2% during setup) was used while positioning the rat into a custom-made MRI cradle. Electrical stimulation was enabled using two pairs of stainless-steel electrodes inserted in the rat forepaws, between digits 2 and 3, and digits 4 and 5, respectively. A catheter was inserted subcutaneously on the back of the animal for medetomidine administration. Once the setup was finalized, an initial bolus of 0.1 mg/kg of medetomidine was injected, while gradually reducing the isoflurane anesthesia. Ten minutes later the isoflurane was completely interrupted, and another five minutes later a continuous medetomidine infusion at 0.1 mg/kg/h was started. The total sedation time under medetomidine was around two hours and a half, with an average respiration rate of 67 ± 18 breaths per minute and an average temperature of 38 ± 0.5°C across all rats and acquisitions. An oxygen/air supply of 20%/80% was maintained throughout the experiment. At the end of the experiment, animals were woken up by intramuscular injection of atipamezole at 0.5 mg/kg.

### 2.2 Paradigm Design

A custom triggering routine making use of the Transistor-Transistor Logic (TTL) pulse output delivered by the MRI scanner was implemented to generate a block paradigm for the dMRI and BOLD-fMRI acquisitions with alternating cycles of 28 s of rest and 28 s of stimulation. The stimulation consisted of 0.3 ms pulses at 2 mA and 8 Hz frequency and was delivered by an A365 stimulus isolator interfaced with a DS8000 digital stimulator (World Precision Instruments, Stevenage, UK). Twenty-eight images for each rest and stimulus intervals were acquired during the BOLD-fMRI (TR = 1 s), and fourteen during the dMRI (TR = 2 s) block designs, respectively.

### 2.3 Data Acquisition

**Figure 1** details the experimental design. The acquisition of functional runs comprising both dMRI and BOLD-fMRI sequences started 30 minutes after isoflurane cessation (**Figure 1A**). During that 30-minute wait time for isoflurane clearance, a B_0_ map for field-map based shimming of the rat brain with default acquisition parameters and a T_2_-weighted (T_2_w) structural acquisition for anatomical reference were performed. The T_2_w structural scan was acquired using a 2D multi-slice RARE sequence, with the following parameters: TE = 6 ms, TR = 2.5 s, matrix size 160 × 160 x 45, in-plane resolution 125 x 125 μm^2^, slice thickness 0.5 mm and RARE factor = 1.

**Figure 1:**
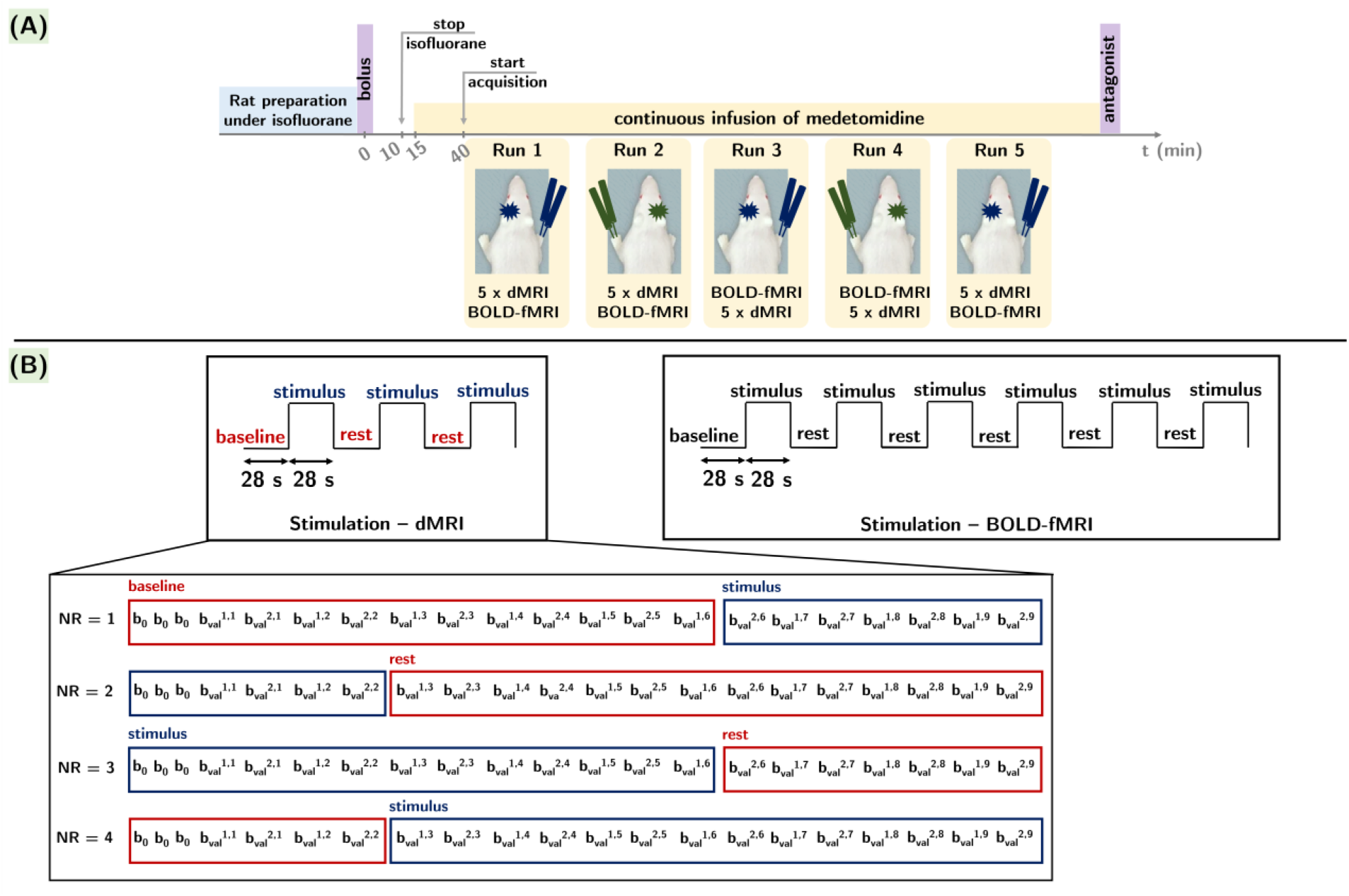
Data acquisition protocol – (A): Experimental timeline. (B): Each functional run included five different diffusion sequences with varying diffusion time values and one BOLD-fMRI sequence. For each diffusion time, the timing of the three epochs of alternating 28 s rest and 28 s stimulus intervals was set such as to acquire NR = 2 repetitions of 21 measurements – nine directions for two b-values (b_val_^i,j^ with i = 1; 2 ms/µm^2^ and j = 9 directions) and three b_0_ acquisitions for each condition (rest vs. stimulation). One BOLD-fMRI acquisition included the acquisition of 336 volumes equally spread across six epochs of alternating 28 s rest and 28 s stimulus intervals.

Five functional runs were acquired per rat (only four runs for two rats). Every functional run was composed of five dMRI acquisitions with different diffusion times (Δ = 9.5; 15; 20; 25; 30 ms) and one BOLD-fMRI acquisition. From one functional run to another, the forepaw used for stimulation was alternated and within each run, the order of the dMRI/BOLD-fMRI acquisitions was randomized. When tallying the total number of animals and functional runs, 24 dMRI datasets per diffusion time and per stimulated forepaw and 24 BOLD-fMRI time-series per stimulated forepaw were acquired. **Table 1** summarizes the acquisition times for the various sequences in the protocol, as well as the time needed for the acquisition of one functional run and for the entire experimental protocol for one rat.

**Table 1:** Acquisition times for the individual dMRI and BOLD-fMRI sequences, one functional run, the B0 map, the T2w structural sequence and finally, for the entire protocol for one rat. For each functional run, one dMRI and one BOLD-fMRI reverse-phase encode acquisition was performed for spatial distortion correction. The total duration of one functional run corresponds to the acquisition times of five dMRI sequences with different diffusion times, one BOLD-fMRI sequence, and the two reverse-phase encode acquisitions. The duration of the entire protocol for one rat includes the acquisition of the B0 map, the structural T2w image and five functional runs.

| Phase encoding: | Forward | Reverse |
| --- | --- | --- |
| **dMRI acquisition** | 2 min 48 s | 8 s |
| **BOLD-fMRI acquisition** | 5 min 36 s | 10 s |
| **1 run (5xdMRI + fMRI)** | 19 min 54 s |  |
| **B<sub>0</sub> map** | 1 min 20 s |  |
| **T<sub>2w</sub>** | 6 min 40 s |  |
| **1 rat dataset** | 1 h 47 min 30 s |  |

### Diffusion MRI

For each diffusion time, a monopolar Pulsed Gradient Spin Echo sequence with Echo-Planar Imaging read-out (PGSE-EPI) was used to acquire two repetitions of 21 measurements – nine directions for two b-values (b_val_^i,j^ with i = 1; 2 ms/µm^2^ and j = 9 directions) and three b = 0 (b_0_) acquisitions – during each condition (rest vs. stimulus). In total, 84 images were equally distributed between three epochs of alternating rest and stimulus intervals (**Figure 1B**) for a total scan time of 2 min 48 s. The remaining experimental parameters included: gradient pulse duration δ = 4 ms, TE = 48 ms, TR = 2 s, matrix size 80 x 80, 21 slices, in-plane resolution 0.25 x 0.25 µm^2^, slice thickness 0.75 µm and partial Fourier acceleration factor of 1.1 in the phase direction. In a separate scan, three b_0_ images were acquired with reversed EPI phase-encode direction, without stimulation, for spatial distortion correction.

### BOLD-fMRI

Data was acquired using a gradient-echo EPI (GE-EPI) sequence with the following parameters: TE = 11.1 ms, TR = 1 s, and matrix size, number of slices, in-plane resolution and slice thickness matching to the dMRI acquisitions (80 x 80, 21, 250 x 250 µm^2^ and 750 µm, respectively). A partial Fourier acceleration factor of 1.2 was employed for the phase direction. A total number of 336 images were acquired during six epochs of alternating rest and stimulus intervals (**Figure 1B**), resulting in 168 images for each rest and stimulus condition, collected in 5 min 36 s. In a separate scan, ten GE-EPI images were acquired with reversed EPI phase-encode direction, without stimulation, for spatial distortion correction.

### 2.4 Data Processing

#### Diffusion MRI

Pre-processing steps, conducted in python (v.3.9, Python Software Foundation), are illustrated in **Figure 2**. The first step (**Figure 2A**) included denoising (Tournier et al., 2019; Veraart et al., 2016), Gibbs unringing (Kellner et al., 2016; Tournier et al., 2019), topup (Andersson et al., 2003) and eddy (Andersson & Sotiropoulos, 2016) correction for each run separately. A rat brain mask was estimated using the atlasbrex (Lohmeier et al., 2019) routine. The two repetitions of diffusion-weighted volumes and the b_0_ images acquired for each rest and stimulus condition were averaged, and the resulting 19 images per condition were used to perform a voxel-wise estimation (Hansen, Lund, et al., 2016; Hansen, Shemesh, et al., 2016) of MD^rest^, MD^stimulus^, MK^rest^ and MK^stimulus^ maps for each diffusion time using Matlab (v.R2021b, MathWorks Inc.) scripts (https://github.com/sunenj/Fast-diffusion-kurtosis-imaging-DKI). Voxels yielding unphysical values due to noise or partial volume (e.g. negative kurtosis or diffusivity) were excluded from further analysis.

**Figure 2:**
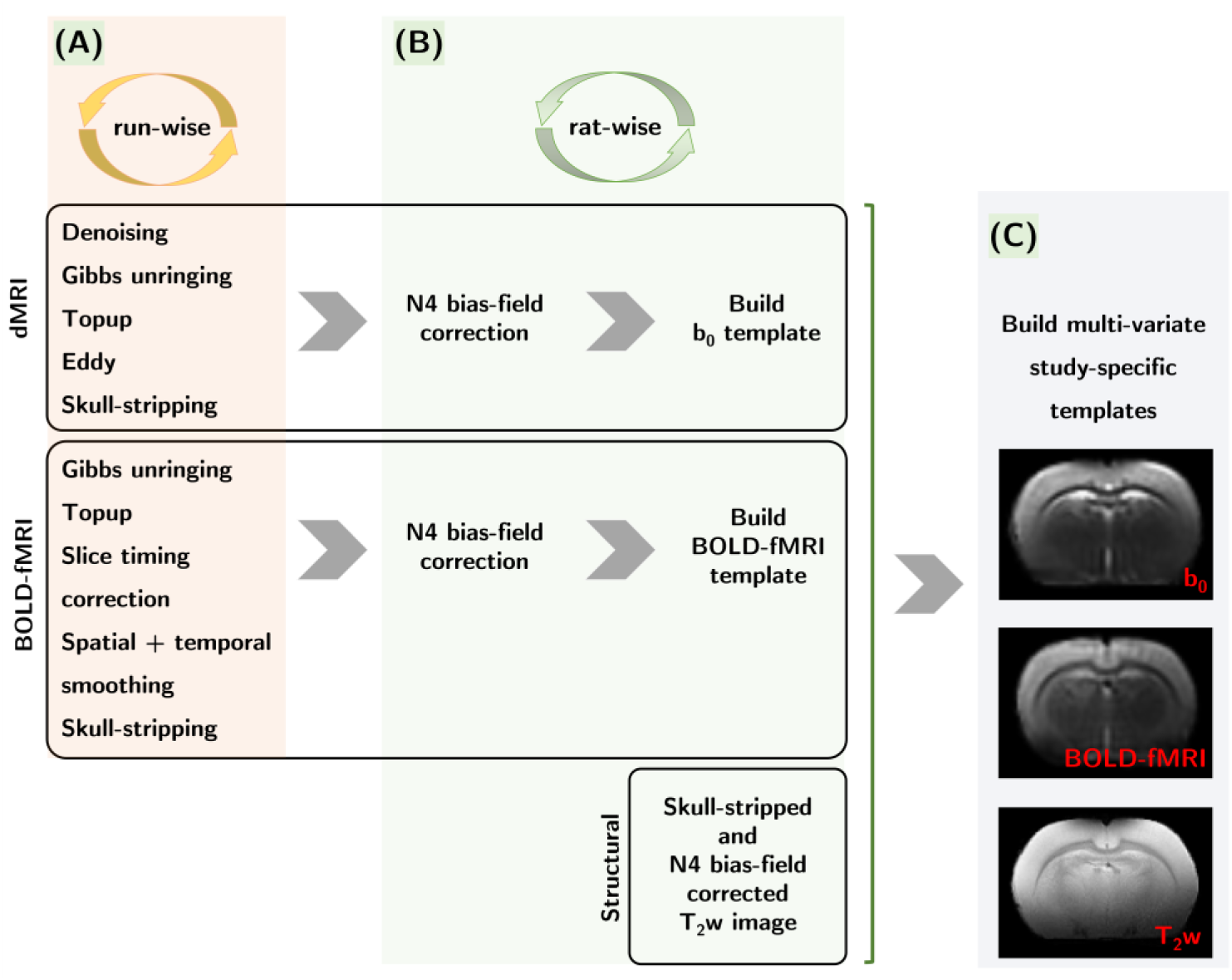
Pre-processing steps – (A): The five diffusion datasets from each functional run were pooled and underwent denoising, Gibbs-unringing, topup and eddy correction, and skull-stripping, while for the BOLD-fMRI time-series topup, slice-timing correction, spatial and temporal smoothing, and skull-stripping were applied for each functional run. (B): Images were then N4 bias-field corrected and the extracted b_0_ images and BOLD-fMRI time-series were used to build corresponding templates for each rat. The skull-stripped and N4 bias-field corrected T_2_w images were jointly used along with the b_0_ and BOLD-fMRI rat-specific templates to construct multi-variate study-specific templates intended to facilitate the propagation of brain segmentations from the WHS T_2_w atlas into the dMRI and BOLD-fMRI individual rat spaces.

#### BOLD-fMRI

Every BOLD-fMRI time-series went through Gibbs unringing (Kellner et al., 2016; Tournier et al., 2019) and topup correction (Andersson et al., 2003) (**Figure 2A**). Prior to the generalized linear model (GLM) analysis in SPM12 (http://www.fil.ion.ucl.ac.uk/spm/software/spm12/), time-series were slice-timing corrected and minimally smoothed (gaussian smoothing kernel with 0.1 x 0.1 x 1 mm^3^ size). High-pass filtering with a cut-off frequency of 0.01 Hz applied on the time-courses ensured the removal of low-frequency noise and drifts. A rat brain mask was estimated using the atlasbrex (Lohmeier et al., 2019) routine for the BOLD-fMRI volumes. For the GLM analysis, a boxcar (Reynaud et al., 2019) response function was used, and statistically significant voxels were detected using a p-value threshold of 0.05 with family-wise error correction. To ensure maximum detection sensitivity, no threshold was applied on the size of the clustered voxels. Prior to ROI-wise quantifications, a brain mask excluding voxels with a signal lower than 10% of the signal measured in the cortex was generated. This step aimed to remove voxels from deeper brain regions where coil sensitivity was heavily attenuated.

### 2.5 Quantification

ANTs (Avants et al., 2009) was employed to generate rat-specific b_0_ and BOLD-fMRI templates from individual images after applying N4 bias-field correction (Tustison et al., 2010) (**Figure 2B**). Individual T_2_w images were also N4 bias-field corrected and skull-stripped and then used jointly along with the b_0_ and BOLD-fMRI rat-specific templates to generate study-specific multivariate b_0_/BOLD-fMRI/T_2_w templates (**Figure 2C**). The WHS atlas (Papp et al., 2014) was registered onto our study-specific T_2_w template and the transformation was applied to the WHS atlas segmentation. The multi-variate templates facilitated the propagation of the brain segmentations into the native dMRI and BOLD-fMRI spaces of each rat.

The segmented regions-of-interest (ROIs) are illustrated in **Figure S1** (Supplementary Materials) and included: primary somatosensory cortex, forelimb area (S1FL), secondary somatosensory cortex (S2), primary motor cortex (M1) as cortical ROIs expected to be involved in the somatosensory processing, thalamus (Tha), striatum (CPu) and hippocampal subfields (Hip) as subcortical ROIs expected to be part of subcortical somatosensory relays, and secondary motor cortex (M2), cingulate cortex (ACC), retrosplenial cortex (RSC) and posterior parietal cortex (PPC) as control cortical ROIs.

Quantifications in each ROI were done in the left hemisphere and were reported as contralateral when the right forepaw was stimulated (expected response in the left S1FL), and ipsilateral when the left forepaw was stimulated (little to no response expected in left S1FL, as the main response should be in the right S1FL).

#### Diffusion MRI

The mean and standard deviation for MD and MK in each ROI were calculated for all diffusion times, runs and rats in the rest and stimulus conditions and were then grouped by stimulated forepaw. The average MD and MK values (± standard error) for each forepaw and each condition were computed to yield MD^rest^(Δ), MD^stimulus^(Δ), MK^rest^(Δ) and MK^stimulus^(Δ) curves in all the investigated brain regions contralaterally and ipsilaterally. MD(Δ) and MK(Δ) time-dependence analyses were conducted in S1FL only. Power-laws for structural disorder (Novikov et al., 2014) and the analytical formulation of time-dependent kurtosis from the two-compartment Kärger model with exchange (Kärger, 1985) were fit to the experimental curves to evaluate potential differences in morphology and permeability between the rest and stimulus conditions. The following equations were used for 1D structural disorder (equations 1 and 3), 2D-3D structural disorder (equations 2 and 4) and the Kärger exchange model (equation 5):

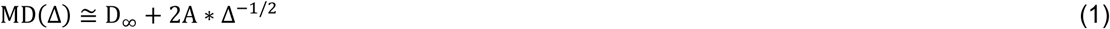

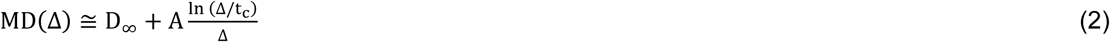

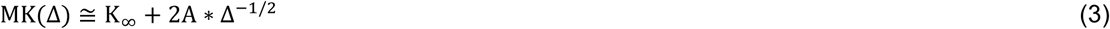

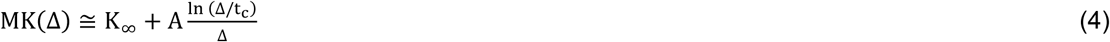

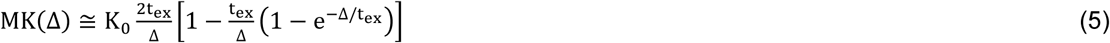

#### BOLD-fMRI

Beyond the GLM analysis, the average and standard deviation of the BOLD-fMRI signal was calculated in each ROI and each volume of the time-series. For each time-series, the BOLD signal in each epoch was normalized by the average signal in the 14 s preceding the stimulation. Then, the BOLD signal response function was calculated by averaging across all epochs, runs and rats. In a separate analysis, and to make a fairer comparison to the MD and MK measures which result from diffusion-weighted images acquired across multiple blocks of rest and stimulation (i.e. yielding one MD and MK estimate per condition), the BOLD signal was also averaged across all timepoints of the rest and stimulus intervals, respectively. As a result, one BOLD amplitude estimate (± standard error) per condition was obtained.

### 2.6 Statistical analyses

All analyses were carried out in RStudio (v.2023.12.1, R Foundation for Statistical Computing), using the lme4 package (Bates et al., 2015). Correction for multiple comparisons using the false discovery rate (FDR) method was applied on each metric (BOLD, MD and MK) separately.

#### Diffusion MRI

The mean MD and MK values quantified in each ROI and for each diffusion time, run, rat and the two conditions were used for the statistical analyses (240 total measurements per ROI). The hierarchical structure of our dMRI data indicated that a mixed effects model would be the most suitable approach. The choice criteria for the statistical model are detailed in **Table S1** (Supplementary Materials). Our factorial design included two main effects: the condition of the rat during the measurement (i.e, rest vs. stimulus) and the diffusion time (five different values) in order to detect statistically significant changes between the rest and stimulus measurements on the one hand, and between measurements at various diffusion times (time-dependence) on the other hand. Two random effects were also considered - the run and the subject indexes. The run index may contribute to random variability in the data due to its temporal spacing in relation to the switch-off of isoflurane anesthesia, slight variations in the temperature and respiration rate across the runs and biological variability in the rat response to the forepaw stimulus. The subject index represents the random variability introduced by biological differences between the rats.

#### BOLD-fMRI

The average BOLD amplitudes estimated for the rest and stimulus conditions were used to determine statistically significant differences in each ROI (48 total measurements per ROI). The statistical design including the state of the rat during the measurement (i.e, rest vs. stimulus) was the most suitable for the evaluation of significant differences as indicated in **Table S2** (Supplementary Materials).

## 3. RESULTS

### 3.1 BOLD-fMRI time-series analyses

A few examples of spatial localization for the statistically significant activation clusters resulting from the GLM analysis conducted on the BOLD-fMRI time-series data corresponding to unilateral right forepaw stimulation are illustrated in **Figure S2** (Supplementary Materials). Globally, the activated area extended over a minimum of five adjacent slices and was mainly localized in the left primary somatosensory cortex. The total number of voxels in the activated region for the 24 datasets (from ten rats) ranged from 14 to 431, with a mean of 179 voxels/cluster. On average, around 80% of the voxels were situated within the primary somatosensory cortex, while the remaining 20% were distributed between the motor cortex (17%) and the secondary somatosensory cortex (3%). Only for five datasets out of twenty-four the activated area presented sparse voxel distributions in the thalamus (< 6%), and less than five datasets presented sparse voxel distributions in the cingulate cortex (<3%), retrosplenial cortex (<3%) auditory cortex (<1.1%), hippocampal subfields (<0.5%) and subicular complex (<0.5%). Percentages are defined with respect to the number of significant voxels detected in the whole brain. The GLM analysis did not unveil any activation clusters in the ipsilateral hemisphere.

### 3.2 Rest vs. stimulus BOLD, MD and MK changes

An example of BOLD image, MD and MK parametric maps for one rat is displayed in **Figure 3** to illustrate the overall quality of our data.

**Figure 3:**
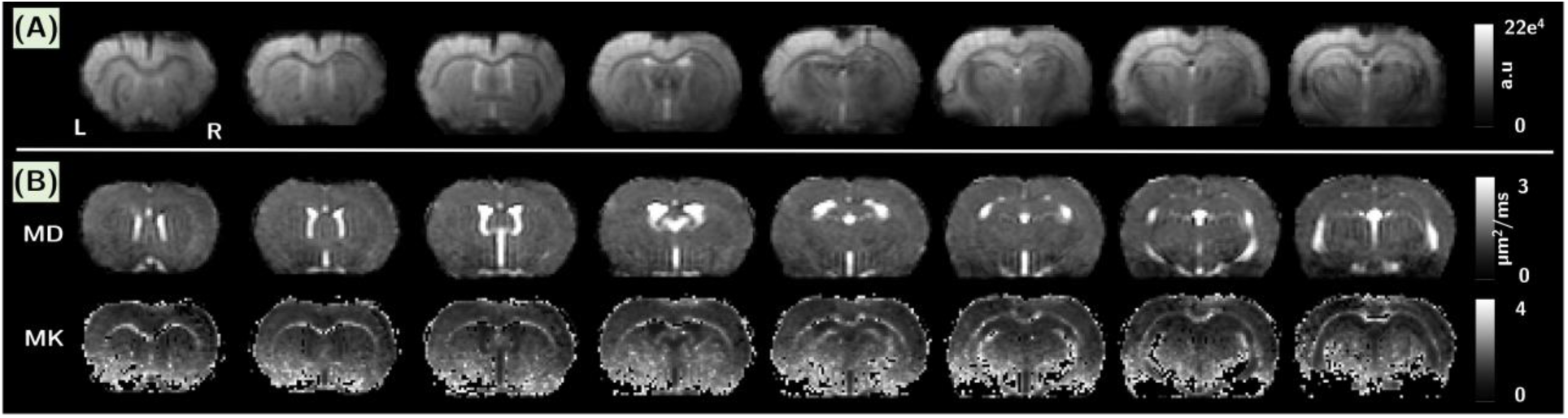
Illustration of the quality of (A): the BOLD-fMRI images after the pre-processing steps and (B): the parametric MD and MK maps, in rostro-caudal slices for one rat. The displayed MD and MK maps correspond to a diffusion time of Δ = 9.5 ms.

In **Table 2** statistically significant amplitude changes in the average BOLD signal, the MD and MK metrics are reported as a percentage of increase or decrease during stimulation with respect to the rest condition. The associated p-values are reported in **Table S3** (Supplementary Materials). We underline that the significant BOLD changes result from the analysis of the normalized time-series signal averaged across each ROI and all rest vs. stimulus timepoints. This approach mirrors the sensitivity of MD and MK estimates, which are also averaged across entire anatomical ROIs and result from diffusion-weighted measurements spread across all the rest vs. stimulus timepoints during each functional acquisition. For MD and MK, the amplitude change was calculated for each diffusion time and only the absolute maximum value was reported. The diffusion times corresponding to the absolute maximum amplitude change are reported in **Table S4** (Supplementary Materials).

**Table 2:** Percentage of signal change during stimulation with respect to the resting condition in the ROIs presenting significant modifications. The BOLD signal changes represent the average relative difference between all rest and stimulus time-points, while the MD and MK changes represent the maximum relative difference between the rest and stimulus conditions obtained across the five diffusion times. The polarity of the signal change is highlighted in colors (green and red for positive and negative changes, respectively). * p < 0.05, ** p < 0.01, *** p < 0.001, with FDR correction for multiple comparisons. The CPu presented a weakly significant BOLD signal decrease (p < 0.05) contralaterally that did not survive correction for multiple comparisons. Globally, the absolute amplitude of the significant MD and MK changes is higher than for BOLD.

| % change |  | S1FL | M1 | S2 | Tha | Hip | CPu |
| --- | --- | --- | --- | --- | --- | --- | --- |
| Contra-lateral | BOLD | +1.2 (***) | +0.2 (*) | +0.3 (**) | - | - | -0.1 |
|  | MD | -1.1 (***) | -1.2 (***) | +1.5 (***) | +1.9 (***) | +1.6 (***) | +1.4 (***) |
|  | MK | -4.9 (***) | - | - | +2.4 (***) | +2.4 (***) | +1.5 (*) |
| Ipsi-lateral | BOLD | - | - | - | - | - | - |
|  | MD | -0.8 (*) | - | +1.6 (***) | +1.6 (***) | +1.3 (***) | +1.6 (***) |
|  | MK | - | - | +3.1 (*) | +1.9 (**) | - | +1.7 (***) |

#### Somatosensory cortical brain regions

The main brain region involved in processing forelimb stimulation – S1FL – presented an average BOLD signal increase of +1.2% contralaterally (**Table 2**, **Figure 4A**). This finding is consistent with the location of the significant signal variations detected at the voxel level in the GLM analysis (**Figure S2,** Supplementary Materials) and the positive response function calculated in the ROI by averaging across all epochs and all rats (**Figure S3A**, Supplementary Materials). Ipsilaterally, BOLD signal changes in whole ROI were not significant, consistent with the response function displayed in **Figure S3A** (Supplementary Materials). On the other hand, diffusion metrics presented a significant decrease in contralateral S1FL during the stimulus condition (**Figures 4B,D**), with maximum amplitudes of -1.1% and -4.9% for MD and MK, respectively (**Table 2**). Ipsilaterally, a weak significant decrease in MD during the stimulation was also found, while no significant changes in MK were detected (**Figures 4C,E**).

**Figure 4:**
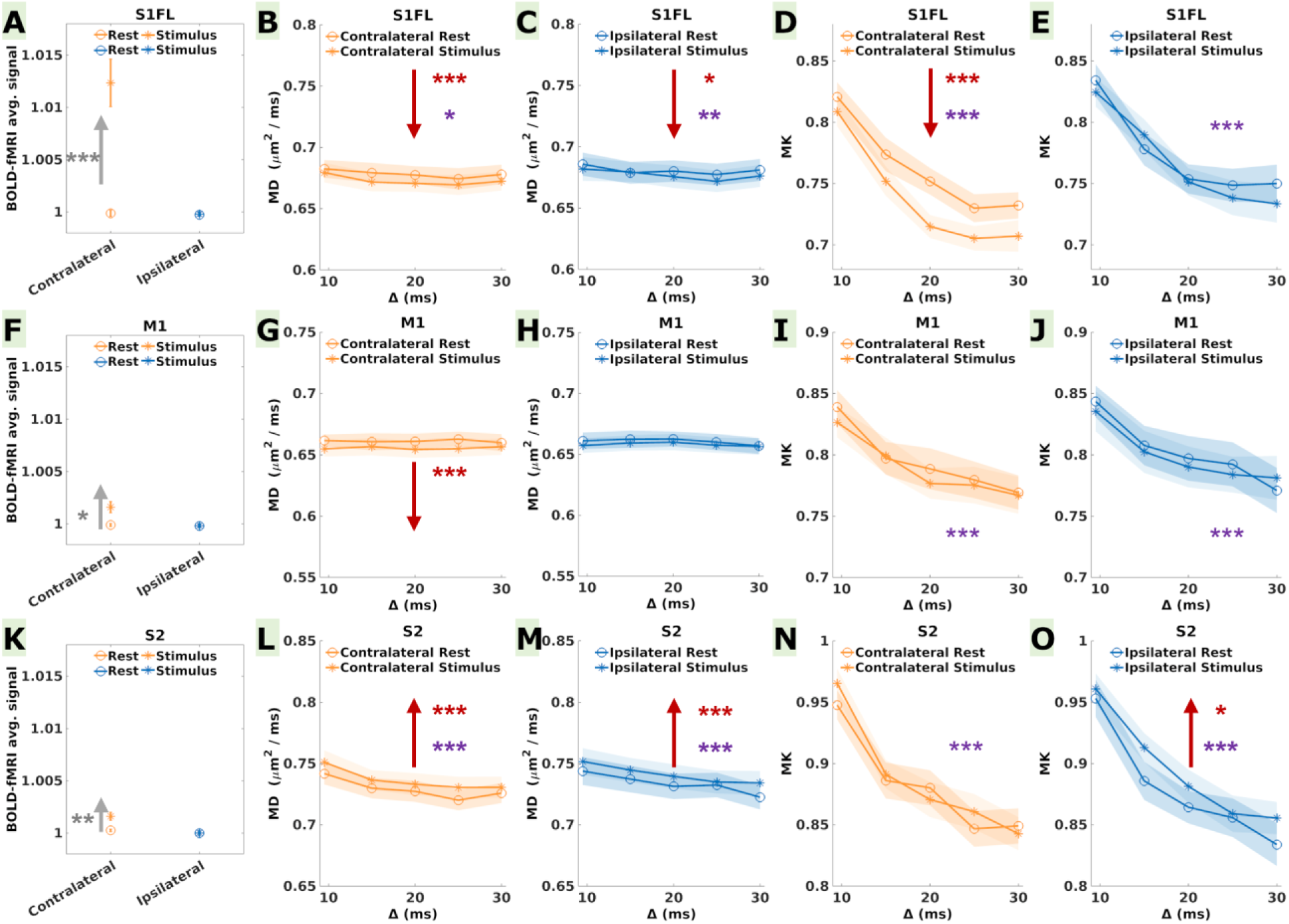
Average BOLD signal, MD and MK during the rest and stimulus conditions in the cortical somatosensory brain regions: (A-E) S1FL, (F-J) M1, and (K-O) S2, contralaterally (blue) and ipsilaterally (orange). For BOLD, statistical differences between the mean across the rest and stimulus time-points are indicated in gray asterisks (the error bars represent the standard error calculated over 24 measurements in each condition). Red and purple asterisks in the MD and MK plots indicate statistically significant differences between the rest and stimulus conditions and between the diffusion time values, respectively (the error bars represent the standard error calculated over 24 measurements in each condition and each diffusion time). P-values are reported as: * p < 0.05, ** p < 0.01, *** p < 0.001 (FDR correction for multiple comparisons). Upwards/downwards red and gray arrows indicate an increase/decrease in the quantified metrics. The significant average BOLD increase in contralateral S1FL was accompanied by a significant decrease in MD and MK, while in contralateral M1 a significant average BOLD increase was paired with a significant decrease in MD only. Ipsilaterally, only MD in S1FL decreased significantly. In contralateral S2 a significant BOLD signal increase was accompanied by a significant increase in MD and MK, contrasting the results in the primary sensorimotor cortices. Ipsilaterally, a significant increase was found only for MK.

In contralateral M1, the average BOLD signal increase (**Figure 4F**) was significant, but lower in amplitude than in S1FL, reaching only +0.2% (**Table 2**). The response function in **Figure S3B** (Supplementary Materials) shows a clear increase in the BOLD signal during the stimulation time-window. Furthermore, a significant MD decrease with an amplitude of -1.2%, remarkably similar in effect size to the MD decrease in S1FL, was measured (**Figure 4G, Table2**). No significant changes were found ipsilaterally (**Figures 4H,J**).

Contralateral S2 also presented an increase in the average BOLD signal during stimulation, with an amplitude of +0.3% (**Figure 4K**, **Table 2**) and a distinctive positive response function (**Figure S3C**, Supplementary Materials). Ipsilaterally, there were no significant BOLD signal variations (**Table 2**). However, a significant increase was found in contralateral S2 for MD (**Figure 4L**), contrasting the trends in S1FL and M1. Remarkably, the MD increase was significant both contra- and ipsilaterally (**Figure 4M**), with maximum amplitudes of +1.5% and +1.6%, respectively (**Table 2**). A significant MK increase of +3.1% was measured only ipsilaterally (**Figure 4O**).

#### Subcortical somatosensory relays

In the Tha and the Hip, no significant changes in the average BOLD signal were found contralaterally and ipsilaterally (**Figures 5A,F**) and the response functions did not display any changes during stimulation (**Figures S3E,F**). Remarkably, contralateral CPu displayed a weak decrease in the average BOLD signal (**Figure 5K**, **Table 2**) with an amplitude of -0.1% and a negative response function (**Figure S3G**, Supplementary Materials). However, the p-value was no longer significant after correction for multiple comparisons.

**Figure 5:**
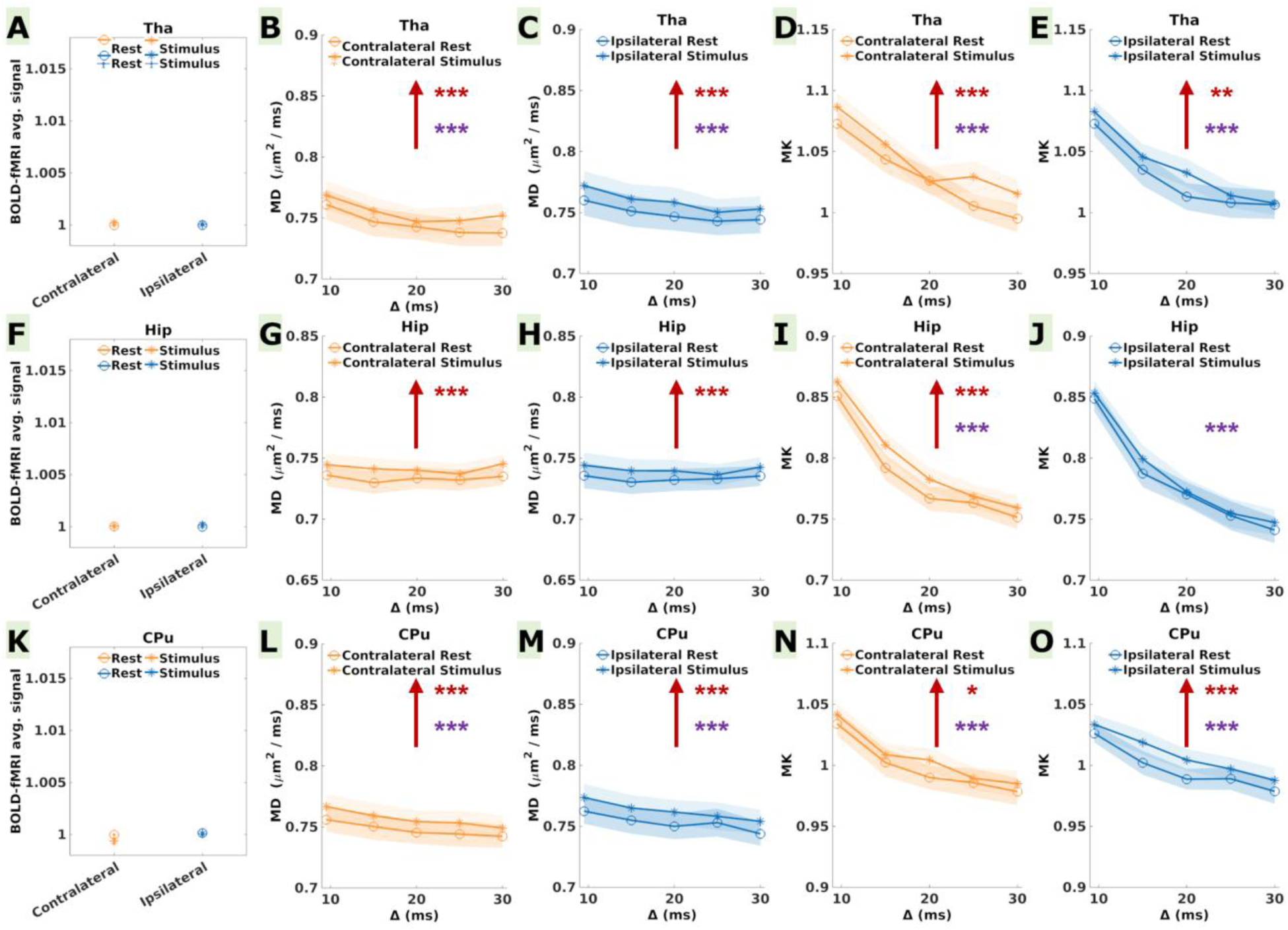
Average BOLD signal, MD and MK during the rest and stimulus conditions in the subcortical somatosensory relays (A-E) Tha, (F-J) Hip and (K-O) CPu, contralaterally (blue) and ipsilaterally (orange). For BOLD, statistical differences between the mean across the rest and stimulus time-points are indicated in gray asterisks (the error bars represent the standard error calculated over 24 measurements in each condition). Red and purple asterisks in the MD and MK plots indicate statistically significant differences between the rest and stimulus conditions and between the diffusion time values, respectively (the error bars represent the standard error calculated over 24 measurements in each condition and each diffusion time). P-values are reported as: * p < 0.05, ** p < 0.01, *** p < 0.001 (FDR correction for multiple comparisons). Upwards/downwards red and gray arrows indicate an increase/decrease in the quantified metrics. Overall, all subcortical ROIs displayed a significant increase in MD and MK during stimulation bilaterally, and only contralateral CPu displayed a very weak decrease in BOLD, however non-significant after FDR correction for multiple comparisons.

Similarly to S2, the subcortical brain regions presented a significant increase in MD both contralaterally and ipsilaterally (**Figures 5B,C,G,H,L and M**), with amplitudes in the range of +1.3 to +1.9%. MK also increased significantly both contralaterally and ipsilaterally in all subcortical ROIs (except for ipsilateral Hip) with amplitudes ranging from +1.5% to +2.4% (**Figures 5D,E,I,J,N and O**).

#### Control cortical brain regions

M2, ACC, RSC and PPC did not display any significant changes in the average BOLD signal, MD or MK between the rest and stimulus conditions (**Figure S4,** Supplementary Materials).

To summarize, in the primary sensorimotor cortices, we report a significant decrease in MD in bilateral S1FL and contralateral M1, and a significant decrease in MK in contralateral S1FL. These changes were paralleled by a significant BOLD signal increase in contralateral S1FL and M1. In the secondary somatosensory cortex and subcortical regions, we report an increase in MD in bilateral S2, Tha, Hip and CPu, and an increase in MK in bilateral Tha and CPu, ipsilateral S2 and contralateral Hip, while at the BOLD signal level we only found an increase in contralateral S2 and a very mild decrease in contralateral CPu. Globally, significant MD and MK changes were consistently on the order of 1% or higher throughout the investigated brain regions, while the reported significant BOLD signal changes were higher than 1% in S1FL and lower than 0.5% in S2 and M1.

### 3.3 MD and MK time-dependence

Measurable time-dependence was found for MD and MK bilaterally in most brain regions, except for MD in bilateral M1, M2 and Hip. The corresponding p-values are reported in **Table S5** (Supplementary Materials).

#### S1FL time-dependence analysis

Analysis of the corrected Akaike information criterion (AICc) values (**Table 3**) associated with fitting structural disorder models (equations 1 and 2) to the MD(Δ) decay in S1FL at rest and during stimulation (**Figures 6A,B,C,D**), revealed a superior quality of fit for the 1D structural disorder model. Indeed, we report lower AICc values along with greater precision in the estimated coefficients. Consequently, the underlying mechanisms behind MD(Δ) time-dependence align more closely with a description based on 1D structural irregularities encountered by water molecules within cell processes, rather than one based on the 2D-3D geometrical details of the extra-cellular space. Consistency in the fitted A and D_∞_ coefficients across conditions (contralateral/ipsilateral, rest/stimulation) indicates that the signature of 1D structural disorder was stable, proffering the hypothesis that differences in MD between rest and stimulation might not arise from alterations in the one-dimensional tissue organization.

**Figure 6:**
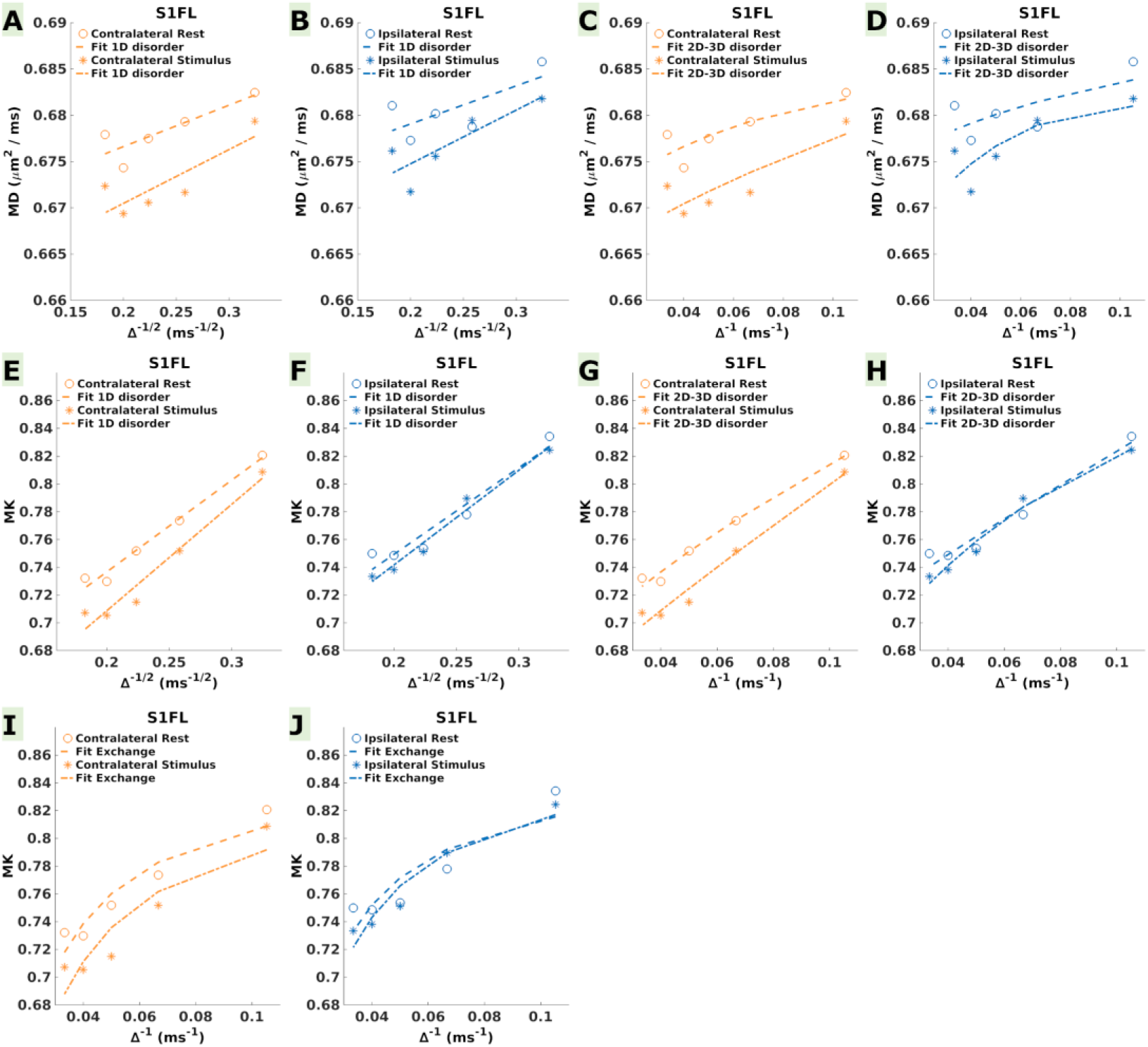
Power-law fit curves (dotted lines) in S1FL for (A,B) structural 1D disorder to MD time-dependence, (C,D) structural 2D-3D disorder to MD time-dependence, (E,F) structural 1D disorder to MK time-dependence, (G,H) structural 2D-3D disorder to MK time-dependence), and (I,J) the Kärger model kurtosis to MK time-dependence.

**Table 3:** Results of the 1D and 2D-3D structural disorder models fit on the MD time-dependence. As evidenced by the lower AICc values and greater precision in the estimated coefficients, the 1D structural disorder model has a superior fit quality for the MD time-dependent data. The minimal differences in AICc, A, and D_∞_ between the rest and stimulus conditions suggest that the characteristics of one-dimensional structural disorder remain consistent across both states, implying that the one-dimensional tissue organization changes might not be responsible for the MD drop during stimulation.

| MD | 1D disorder |  | 2D-3D disorder |  |  |
| --- | --- | --- | --- | --- | --- |
| | A | $D_{\infty}$ | A | $t_c$ (ms) | $D_{\infty}$ |
| Contralateral Rest | $0.02 \pm 0.008$ | $0.67 \pm 0.004$ | $0.08 \pm 0.26$ | $2.03 \pm 6.4$ | $0.67 \pm 0.02$ |
|  | AICc = -42 |  | AIC = -21 |  |  |
| Contralateral Stimulus | $0.03 \pm 0.01$ | $0.66 \pm 0.005$ | $0.04 \pm 0.33$ | $0.29 \pm 7.1$ | $0.66 \pm 0.02$ |
|  | AICc = -38 |  | AIC = -19 |  |  |
| Ipsilateral Rest | $0.02 \pm 0.01$ | $0.67 \pm 0.006$ | $0.05 \pm 0.37$ | $1.37 \pm 13.1$ | $0.67 \pm 0.02$ |
|  | AICc = -38 |  | AIC = -18 |  |  |
| Ipsilateral Stimulus | $0.03 \pm 0.01$ | $0.66 \pm 0.005$ | $0.22 \pm 0.36$ | $3.37 \pm 2.8$ | $0.66 \pm 0.02$ |
|  | AICc = -39 |  | AIC = -18 |  |  |

Upon analyzing MK(Δ) decay (equations 3 and 4), the 1D structural disorder model demonstrated once again to be better suited. This consistency in model preference across MD and MK reinforces the robustness of the 1D structural disorder concept in characterizing the underlying tissue microstructural features. The higher AICc values obtained from fitting the structural disorder models to MK(Δ) (**Figures 6E-H**, **Table 4**) compared to those for MD(Δ) underscore the complexity of MK time-dependence, which is influenced by additional factors, such as exchange. Indeed, we found comparable AICc values between the 1D structural disorder and the Kärger exchange models (**Figures 6E,F,I,J**, **Table 4**), reinforcing the notion that the underlying mechanisms for both models serve as confounding factors affecting the MK time-dependence. The Kärger model fit to MK(Δ) (equation 5) yielded a shorter exchange time during stimulation (t_ex_^stimulus^ = 45.1 ± 12.0 ms vs. t_ex_^rest^ = 53.8 ± 11.1 ms contralaterally, and t_ex_^stimulus^ = 51.5 ± 8.8 ms vs. t_ex_^rest^ = 60.7 ± 19.5 ms ipsilaterally), suggesting increased membrane permeability during the stimulus condition. However, the difference in exchange time during rest and stimulation fell within the range of the estimation uncertainty (**Table 4**), possibly reflecting the complex interplay between structural disorder and exchange mechanisms.

**Table 4:**
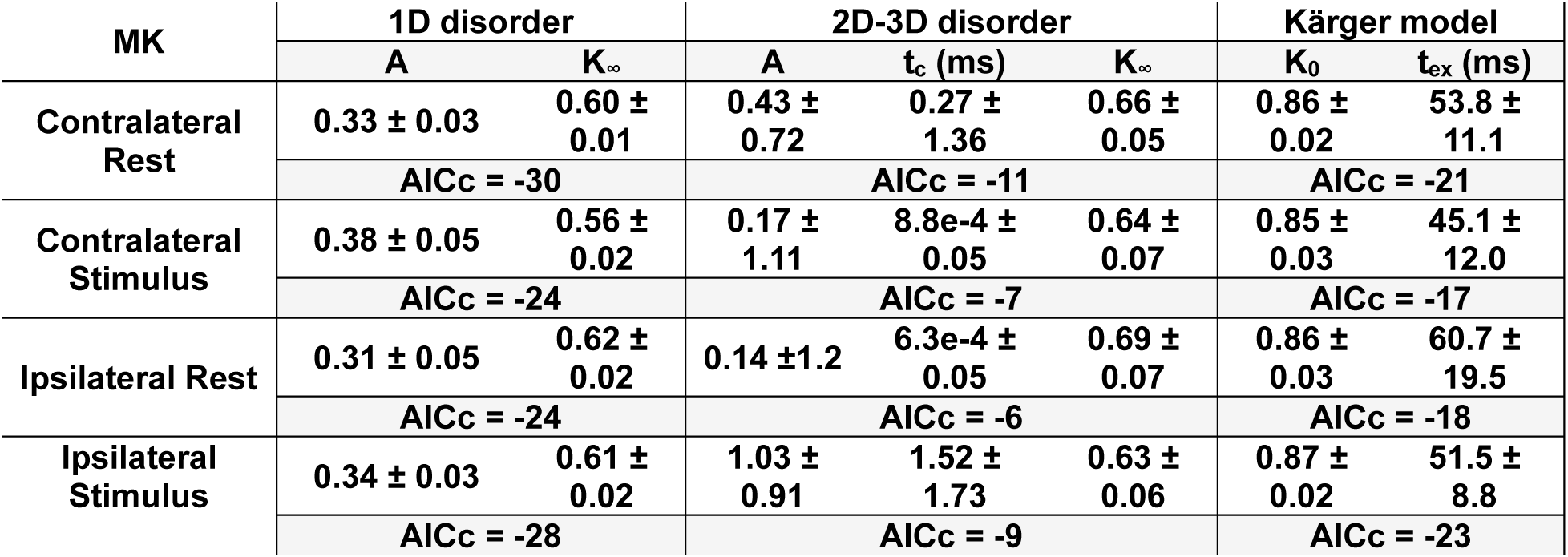
Results of the 1D and 2D-3D structural disorders, and Kärger model kurtosis fit on the MK time-dependence. Consistent with the findings for MD time-dependence, the 1D structural disorder model exhibited lower AICc values and greater precision in the estimated coefficients compared to the 2D-3D structural disorder fit. Both structural disorder and exchange mechanisms influence MK time-dependence, which is evident in the comparable predictive abilities of the 1D structural disorder and Karger models when applied to the MK(Δ) data. Despite the intricate interplay between structural disorder and exchange, a slight decrease in exchange time was observed during stimulation.

## 4. DISCUSSION

In this paper, we report brain region-specific changes in MD and MK related to a task of forepaw electrical stimulation. In particular, we present the first evidence of a decrease in MK, paired with previously reported decreased MD (Darquié et al., 2001; Yacoub et al., 2008), associated with neuronal activity within the primary somatosensory cortex in the rodent brain, and an increase in MD and MK in the secondary somatosensory and subcortical areas. These modifications did not parallel changes in the BOLD signal in a systematic way, pointing to a disconnection of underlying functional contrast sources between diffusion weighting and BOLD.

### 4.1 Somatosensory cortical and subcortical pathways

While the primary somatosensory cortex is generally the main focus in functional studies of forepaw or hindpaw electrostimulation (Bock et al., 1998; Brinker et al., 1999; Spenger et al., 2000; Masamoto et al., 2006; Pelled et al., 2009; T. Kim et al., 2010; Adamczak et al., 2010; Goloshevsky et al., 2011; Baltes et al., 2011), a detectable BOLD response was previously reported (Keilholz et al., 2004; Weber et al., 2006; Cho et al., 2007; Zhao et al., 2008; Bosshard et al., 2010; Boussida et al., 2017; Jung et al., 2019, 2021) in the sensorimotor and subcortical areas found to present task-induced MD and MK changes in the current study. In addition to activation in the contralateral S1FL, a weak BOLD response was also formerly detected ipsilaterally (Boussida et al., 2017; Jung et al., 2019) and was attributed to somatosensory activity via inter-hemispheric sensory pathways (Jung et al., 2019). Furthermore, a positive BOLD response in the contralateral M1 was found in several other studies (Weber et al., 2006; Cho et al., 2007; Bosshard et al., 2010; Boussida et al., 2017; Jung et al., 2019, 2021), although it was construed as an overflow in some cases (Weber et al., 2006) or an antidromic stimulation of efferent motor fibers in others (Bosshard et al., 2010; Cho et al., 2007). Nevertheless, studies using retrograde tracing techniques have clearly demonstrated the existence of cortical projections to M1, mainly originating from the primary somatosensory region in the same hemisphere (Colechio & Alloway, 2009; Yamawaki et al., 2021). Moreover, the primary somatosensory cortex is part of a complex intra-cortical and cortico-thalamo-cortical network which facilitates the flow of sensory information (Jabaudon, 2017). Another retrograde tracer injection study revealed the existence of thalamocortical projections ascending through the caudate putamen or directly from the ventral posterolateral thalamic nucleus to the primary and secondary somatosensory cortices, while the latter cortices were shown to be reciprocally connected (Liao & Yen, 2008). Cortical afferents from S1 and M1 to the CPu were suggested as mediators for sensorimotor integration in the basal ganglia using tracer anterograde labeling (Ramanathan et al., 2002), while somatosensory processing associated with a task of whisker pad stimulation was shown to trigger hippocampal responses in electrophysiological recordings (Bellistri et al., 2013). From the perspective of BOLD, positive responses were measured in S2 (Bosshard et al., 2010; Boussida et al., 2017; Jung et al., 2019, 2021; Keilholz et al., 2004; Weber et al., 2006; Zhao et al., 2008) and Tha contralaterally (Jung et al., 2019, 2021; Zhao et al., 2008) or bilaterally (Bosshard et al., 2010; Boussida et al., 2017; Keilholz et al., 2004), whereas a negative BOLD response was reported bilaterally in the CPu (Boussida et al., 2017; Zhao et al., 2008). However, none of these studies reported a BOLD response in the hippocampal subfields. Globally, the entirety of this body of literature strongly indicates that brain activity during somatosensory processing extends well beyond S1FL and is relevantly captured by MD and/or MK changes in our study.

### 4.2 Microstructural dynamics underlying MD and MK changes in S1FL and M1

The observed MD and MK changes we report reflect the collective impact of numerous microscopic processes (e.g., geometric changes in the intra- and extracellular spaces resulting from cellular swelling, increased tortuosity and restrictions in the extracellular space, decreased restrictions in the intracellular space, decreased transmembrane permeability, etc.) occurring simultaneously across multiple spatial scales.

The measured MD drop in S1FL and M1 during the stimulus condition was previously reported in human (Darquié et al., 2001; Le Bihan et al., 2006) and rodent (Abe, Tsurugizawa, et al., 2017; Tsurugizawa et al., 2013) studies, and was attributed to cellular swelling and increased tortuosity in the extracellular space. While the precise mechanisms underlying an MD reduction during neural activity remain unclear, functional- induced alterations in water diffusivity may exhibit varying patterns contingent upon the cellular structure(s) or the spatial scale under investigation. For example, in the *Aplysia Californica* buccal ganglia, water diffusivity was reported to increase inside neuronal soma and decrease at the level of the whole tissue upon cellular swelling (Jelescu et al., 2014; Abe, Van Nguyen, et al., 2017), while a closer look at water diffusivity in the synaptic microenvironment revealed a tenfold increase near the excitatory synapses, in contrast to regions farther away from the synaptic site (Paviolo et al., 2022). In our study, time-dependence analyses of MD(Δ) were expected to bring more insight into the intracellular vs. extracellular contributions behind the stimulus-induced MD changes, but the power-law fit for structural disorder did not allow us to unambiguously disentangle between these possible contributions. A comparison of AICc values between the structural disorder models for each condition (rest vs. stimulation) indicates that the 1D structural model provides a better fit for the MD(Δ) data in both rest and stimulation states (**Table 3**). In other words, the intracellular restrictions arising from variations along the main axis of axons or cell processes such as caliber modifications and beadings seem to explain better the MD time-dependence in our data than the extracellular hindrances generated by complex geometrical features such as undulations, branching or interactions between various cellular components in the 3D space. However, the similar microstructural coefficients (A and D_∞_) obtained from fitting the 1D model to the MD time-dependent data at rest and during stimulation suggest that changes in structural disorder within the intracellular compartment may not be the main factor driving the observed MD(Δ) decrease during stimulation. Conversely, the 2D-3D structural disorder also provides comparable AICc values between the rest and the stimulus conditions (**Table 3**), though certain estimated microstructural coefficients (t_c_ and A) differ by up to an order of magnitude between rest and stimulation. Nonetheless, the large uncertainties associated with these parameter estimations limit our ability to draw definitive conclusions or make further inferences about the precise nature of structural modification associated with MD/MK differences between rest and stimulation. Both types of disorder may simultaneously influence the observed MD(Δ) trends, but the precise extent of their contribution remains undetermined.

As for MK, our initial hypothesis posited that it could capture the alterations in trans-membrane permeability linked to the regulation of imbalances induced by physiological and biochemical processes accompanying action potential propagation and synaptic transmission (Bai et al., 2019; Kitaura et al., 2009; Bai et al., 2018; Bellot-Saez et al., 2017; Didier et al., 2018). Notably, the genesis of advanced diffusion biophysical models for gray matter tissues such as Neurite Exchange Imaging (NEXI) (Jelescu et al., 2022; Olesen et al., 2022; Uhl et al., 2024) was prompted by the imperative to address the impact of exchange between the intra- and extracellular compartments on diffusion-weighted signals. Consequently, the imprint of exchange in gray matter is ingrained within the dMRI signal. As such, the measured MK reduction in contralateral S1FL suggested an increase in trans-membrane permeability resulting from reduced water compartmentalization, further confirmed by the slight decrease in the estimated exchange time from MK(Δ) time-dependent analyses. However, MD time-dependence in our data is a strong indicator of the non-gaussian diffusion of water molecules generated by structural irregularities (Novikov et al., 2014). Comparisons between the fit results of structural disorder models to the MD(Δ) data (**Table 3**) show that the 1D model provides a superior fit, along with more precise microstructural estimates. This signature is necessarily mirrored in the MK time-dependence. Thus, the interpretation of MK(Δ) is complicated by the simultaneous contribution of both 1D structural disorder and exchange mechanisms, which require more advanced models and extensive datasets to separate these effects Exchange (Novikov et al., 2023).

### 4.3 Microstructural dynamics underlying MD and MK changes in S2 and subcortical areas

Interestingly, findings in S1FL and M1 were contrasted by the increase in MD and MK in S2 and the subcortical ROIs, possibly reflecting an opposite signature in terms of water mobility across the microstructure, e.g. a different excitatory/inhibitory balance, for the latter brain regions. Indeed, sensorimotor signals were shown to initiate both excitation and inhibition processes in the various hippocampal subfields (Bellistri et al., 2013), an inhibitory postsynaptic potential was reported in various thalamic nuclei (O’Reilly et al., 2021), while the striatum is one of the main hubs for inhibitory interneurons (Fahn et al., 2011; Ramanathan et al., 2002). S2 may not be directly associated with an inhibitory response, but projections from both thalamic nuclei and S1 to S2 might result in both inhibitory and excitatory responses. As for the underlying mechanisms explaining a dominant MD increase during inhibitory/excitatory balance, cellular shrinkage associated with neuronal hyperpolarization (Fraser & Huang, 2004) is one possible hypothesis, while reduced trans-membrane water transport in the hyperpolarized state could be at the origin of an increase in MK. However, as suggested by a recent study focusing on the rat and human striatum (Cerri et al., 2024), the polarity of the functional response as measured by BOLD cannot be solely explained by neuronal activity subcortically, i.e. positive BOLD = excitatory activity and negative BOLD = inhibitory activity, but rather by complex neurochemical feed- forward mechanisms. A different neurochemical environment across the various brain regions involved in somatosensory processing and integration could also have an impact on water diffusivity, and thus lead to the observed MD and MK trends. An alternative hypothesis would involve a region-dependent shift in the dominant mechanisms or water compartmentalization influencing MD and MK. Notably, astrocytes behave as excellent osmometers (Hellas & Andrew, 2021) and play an important role in brain excitability (Schwartzkroin et al., 1998). The astrocytic density is higher in subcortical regions than in the cerebral cortex in the adult rat brain (Savchenko et al., 2000) and changes in astrocyte volume were shown to be dependent on the degree of hypoosmolality of their surrounding medium (Lafrenaye & Simard, 2019), with a direct link between astrocyte volume regulation and membrane permeability (Solenov et al., 2004). Nonetheless, additional studies are warranted to elucidate the precise mechanism at play and their specific relationship with MD and MK.

### 4.4 Dissociation between BOLD and diffusion changes and their respective sensitivities

As previously stated, MD and MK changes during stimulation did not parallel BOLD signal modifications in a systematic way. In contralateral S1FL and M1, the average BOLD signal increase was paired by a decrease in MK and/or MD, while ipsilateral S1FL presented a decrease in MD in the absence of detectable BOLD signal variations. On the other hand, a weak negative BOLD signal was accompanied by strongly significant elevations of MD and MK in the CPu. In the remaining brain regions, the trends were dissociated: the BOLD signal and MD increased in contralateral S2, while no significant BOLD change were found in ipsilateral S2, ipsilateral CPu, bilateral Tha or Hip in the presence of an increase in MD and MK. The absolute amplitude of the BOLD signal changes scaled with the expected functional involvement of each area, with the highest response in S1FL (1.2%) and gradually less in S2 (0.3%), M1 (0.2%) and CPu (0.1%), although whether the order between S2, M1 and CPu reflects the actual strength of functional activity is not known. Intriguingly, the absolute amplitude of the significant MD and MK changes was overall above 1%, except for MD in the ipsilateral S1FL. The absolute maximum amplitude changes for MD were weaker in S1FL and M1 (1.1%/1.2%) than in areas of MD increase, such as contralateral S2 or Tha (1.5%/1.9%). On the other hand, contralateral S1FL showed a marked absolute MK variation (maximum of 4.9%) while the absolute amplitude of the significant changes in Tha and Hip was reduced by half (maximum of 2.4%). While contralateral S1FL is undoubtedly the area of strongest excitatory activity in this task, it remains to be established if the comparable MD amplitude changes between S1FL and the subcortical regions are the result of competing mechanisms that contribute to either an increase or decrease in MD, or a possibly strong inhibitory response in subcortical regions.

Measuring significant diffusion changes below the BOLD detection threshold indicates a higher sensitivity for MD and MK to brain activity, although it is possible that the neurovascular responses in the subcortical areas were too weak to be detected with our experimental setup. Indeed, the acquired dMRI and BOLD-fMRI images reflect the sensitivity profile of a surface coil, with reduced SNR in the deeper brain regions as compared to the cortex. However, both GE-EPI and PGSE-EPI were affected by this profile, with the spin-echo diffusion images possibly even more affected due to larger flip angles and relative inefficiency of the refocusing pulse far from the coil. Furthermore, the GE-EPI sequence was optimized for BOLD sensitivity, following previously established and recommended protocols (Grandjean et al., 2023; Reynaud et al., 2019). Beyond GLM voxel-wise analyses, we attempted to match the sensitivity of BOLD and MD/MK by averaging the BOLD signal across all rest and stimulus intervals for each anatomical ROI. In spite of this, we report an increase in MD and MK in the thalamus and the hippocampus, in the absence of a notable BOLD signal change.

Furthermore, changes in the diffusion metrics remained specific to regions involved in the somatosensory processing and integration, as no MD or MK changes were observed in cortical regions such as ACC, RSC, PPC or M2 which were also devoid of a BOLD response.

### 4.5 Limitations - dMRI

The fast kurtosis approach (Hansen, Lund, et al., 2016) used to estimate MD and MK in a scan time compatible with a functional block design is certainly associated with higher estimation uncertainty as compared to the full kurtosis tensor estimation. In practice, a ‘classical’ protocol for diffusion and kurtosis tensor estimation comprises approximately 30 directions per b-value, well superior to the nine directions per b-value acquired in this study. In addition, the higher intrinsic sensitivity to noise for the kurtosis metrics led to a relatively large MK uncertainty, which also translated into fewer ROIs displaying a significant change in MK as compared to MD. Thus, the analysis of time-dependent MK(Δ) was likely confounded not just by concomitant contributions between structural disorder and exchange, but also by the limited precision of fast MK estimates. Ideally, future studies would employ microstructural biophysical models, such as NEXI (Jelescu et al., 2022; Olesen et al., 2022; Uhl et al., 2024), or more complex models simultaneously accounting for both structural disorder and exchange mechanisms (Novikov et al., 2023), to enable access to compartment-specific water diffusivity changes associated with neural activity. However, the use of multiple dMRI measurements to estimate the parameters of interest limits this approach to task fMRI with long block designs (56 s per epoch in our case, 3 epochs per run) producing a sustained activity over a sufficiently long duration. Such limitations restrict our ability to resolve rapid microstructural changes associated with the neural response and distinguish between the various physiological mechanisms underlying the transient water diffusivity changes. Consequently, the observed MD and MK differences between the rest and stimulation states in this study likely reflect more sustained or cumulative microstructural changes. Faster dMRI acquisition schemes, as for example based on isotropic encoding (Nunes et al., 2019, 2021; Spencer et al., 2024) enable higher temporal resolution acquisitions and could be used in future studies, although their unclear definition of the diffusion time pose additional challenges for the time-dependence analyses.

The low number of measurements per functional run for MD and MK estimation also limited the statistical power of our data, as compared to BOLD where many samples are acquired with high temporal resolution. Therefore, the voxel-wise GLM analysis typical for BOLD-fMRI did not yield any significant results for MD or MK. As a result, we compensated for the low number of available measurements by averaging over anatomical ROIs. The voxel-wise b = 0 SNR ranged from 10 (thalamus) to 20 (S1FL), which translates into good accuracy and precision of MD and MK estimates (see Fig. 2 in (Hansen, Lund, et al., 2016)) in all our ROIs, except thalamus. The ROI averaging over 100 – 1400 voxels ensured the precision was well improved in all cases. The same ROI averaging for the BOLD signal was performed to make the statistical results from the two contrasts more directly comparable.

Another limitation is the possible BOLD-like contribution to MD and MK estimates, i.e. driven by changes in blood volume and oxygenation that cause susceptibility changes in the surrounding tissues. While working with MD and MK, rather than raw diffusion-weighted signals, substantially reduces T2-weighting effects that could reflect BOLD contrast, it is important to note that our experimental design remains somewhat susceptible to these influences. As illustrated in Figure S3, the BOLD response function is not constant during the stimulation time-window and extends into the rest interval for approximately 4 seconds post-stimulus. The dynamic nature of the BOLD response results in a variable T2-weighting bias across the dMRI measurements. Methods such as the Incomplete Initial Nutation Diffusion Imaging offer a promising approach to separate T2 effects from diffusion measurements measurements (Ianuş & Shemesh, 2018; Nunes, Daniel et al., 2018) and could eliminate potential biases in the quantification of MD and MK that arise from time-varying T2-weighting. Nevertheless, the use of an ultra-high magnetic field strength (here, 14.1T) results in substantially shorter T2 relaxation times for blood, thus significantly diminishing the direct contribution of intravascular water to the measured diffusion signal. Conversely, the use of higher magnetic field strengths introduces a new set of challenges. In particular, susceptibility-induced background field gradients around blood vessels become much more significant and their interaction with the diffusion-weighting gradients can lead to an increase in apparent diffusivity (Pampel et al., 2010). We note however that in the case of positive BOLD, we measured a decrease in MD rather than an increase in MD, and we also report MD and MK changes in areas where a measurable BOLD response was absent. The proof that task-driven MD and MK changes are not dependent on vascular response but are instead associated with changes in cellular microstructure remains valid.

Finally, we emphasize that reporting the largest amplitude change in MD and MK across the various diffusion times allowed us to identify the optimum diffusion time value for maximum sensitivity to region-specific functional-induced changes. Overall, the optimal diffusion time should be tuned to the characteristic exchange time of the system. Consequently, it was not surprising that we observed the majority of maximum amplitude changes in MD and MK within the range of 15-25 ms (Jelescu et al., 2022). For whole brain studies at a single diffusion time, a value of around 15 ms represents the best compromise in terms of sensitivity to functional MD and MK changes across brain regions. However, various types of stimuli, differences in the tissue type investigated (White Matter vs. Gray Matter), or the presence of a pathological state might prompt the need for a different acquisition protocol to enhance the functional sensitivity of MD/MK.

### 4.6 Limitations – BOLD-fMRI

Factors such as magnetic field strength (Gati et al., 1997), echo time (Grüne et al., 1999) and temporal SNR (tSNR) (Murphy et al., 2007) all influence BOLD sensitivity. In this study, we leveraged the advantages of an ultra-high magnetic field system to enhance BOLD sensitivity, and the echo time was optimally set according to a recently published consensus protocol for functional connectivity analysis in the rat brain (Grandjean et al., 2023). Nonetheless, lower T_2_* in subcortical brain regions (Boussida et al., 2017) resulting from differences in vascular organization (Sloan et al., 2010) and susceptibility effects as compared to cortical areas, might prompt the need for a different echo time in order to maximize BOLD sensitivity subcortically. The use of a surface transceiver RF coil in our study led to a notable tSNR reduction in the deeper structures of the brain, possibly obscuring weaker BOLD changes. However, the same penalty applied for the diffusion-weighted images. New approaches for the analysis of BOLD-fMRI time-series employing response function tailored (Gonzalez-Castillo et al., 2012; Schilling et al., 2018) by region, task and/or subject could be used in the future for a more robust detection of voxel-wise significant BOLD signal changes beyond the main activated brain regions.

## 5. CONCLUSIONS

To conclude, in this study we have shown an MD and MK reduction during brain activity in contralateral S1FL consistent with changes in cellular morphology and water transport mechanisms, paired with a positive BOLD response. As the MD and MK changes between the rest and stimulus conditions did not parallel the BOLD signal dynamics in a systematic way in all the investigated brain regions, the current study demonstrates the potential of MD and MK to provide complementary functional information to BOLD across multiple cortical and subcortical areas in the rat brain. Remarkably, MD and MK detected functional- induced changes in areas reported in the literature as part of the somatosensory processing and integration pathways, with overall higher sensitivity than BOLD. Future research endeavors will focus on a deeper investigation of the underlying mechanisms behind the elevation in MD and MK associated with neural activity, particularly as observed in S2 and subcortically. Calcium imaging, along with chemogenetic and optogenetic neuromodulation techniques, enable precise targeting of specific cell types in vivo, facilitating the assessment of contributions from various populations, including glial cells and excitatory and inhibitory neurons (Kim & Jun, 2013; Markicevic et al., 2021; Sancataldo et al., 2019). When combined with advanced super-resolution techniques that visualize the organization of the extracellular space at the cellular level (Godin et al., 2017; Gwosch et al., 2020; Tønnesen et al., 2018), these methods can provide a more comprehensive understanding of the brain’s microstructure-function relationship during neural activity, particularly in relation to the dMRI measurements.

## DATA AND CODE AVAILABILITY

All data related to this paper will be publicly available on the Zenodo website once the article is accepted (10.5281/zenodo.1109669).

## AUTHORS CONTRIBUTION

AH: conceptualization, data curation, formal analysis, investigation, methodology, visualization, writing – original draft; TP: methodology, writing – review & editing; IOJ: conceptualization, methodology, funding acquisition, project administration, resources, supervision, writing – original draft.

## ACKNOWLEDGEMENTS

The authors thank Lucas Soustelle (Centre de Résonance Magnétique Biologique et Médicale, Marseille, France) for sharing the diffusion sequence with the reversed phase encoding and for help with the implementation of the trigger in the diffusion sequence; Stefan Mitrea, Analina Hausin and Estelle Gerossier for their assistance with animal setup and monitoring; and Bernard Lanz and Claudia Zanella for providing the RF coil. This work took place at the CIBM Center for Biomedical Imaging, a Swiss research center founded and supported by Lausanne University Hospital (CHUV), the University of Lausanne (UNIL), the Swiss Federal Institute of Technology (EPFL), the University of Geneva (UNIGE) and Geneva University Hospital (HUG) and was supported by the Swiss National Science Foundation under Eccellenza grant PCEFP2_194260. We acknowledge the resources and expertise of the CIBM Center for Biomedical Imaging.

## SUPPLEMENTARY MATERIALS

**Figure S1:**
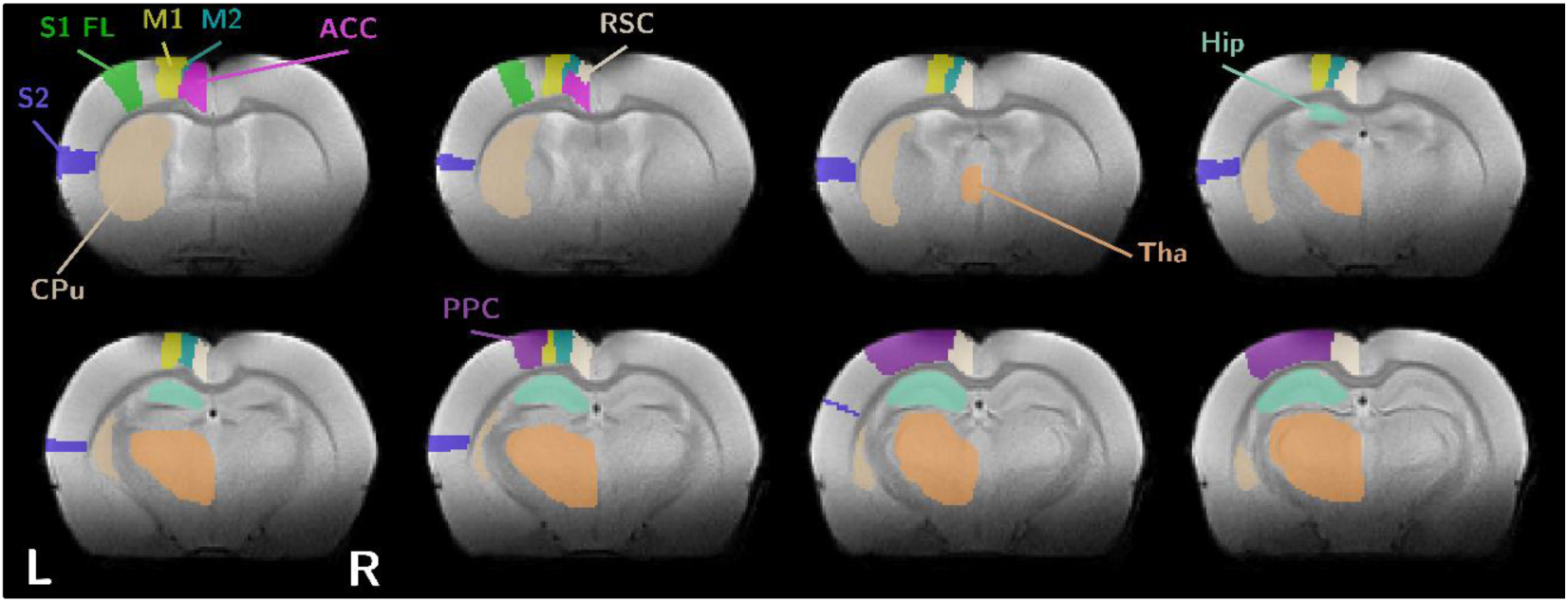
Illustration of the segmentation of various areas investigated in this study on slices spanning from the rostral the caudal brain: primary somatosensory cortex, forelimb area (S1FL), secondary somatosensory cortex (S2), primary motor cortex (M1), secondary motor cortex (M2), cingulate cortex (ACC), retrosplenial cortex (RSC), posterior parietal cortex (PPC), thalamus (Tha), striatum (CPu) and hippocampal subfields (Hip).

**Figure S2:**
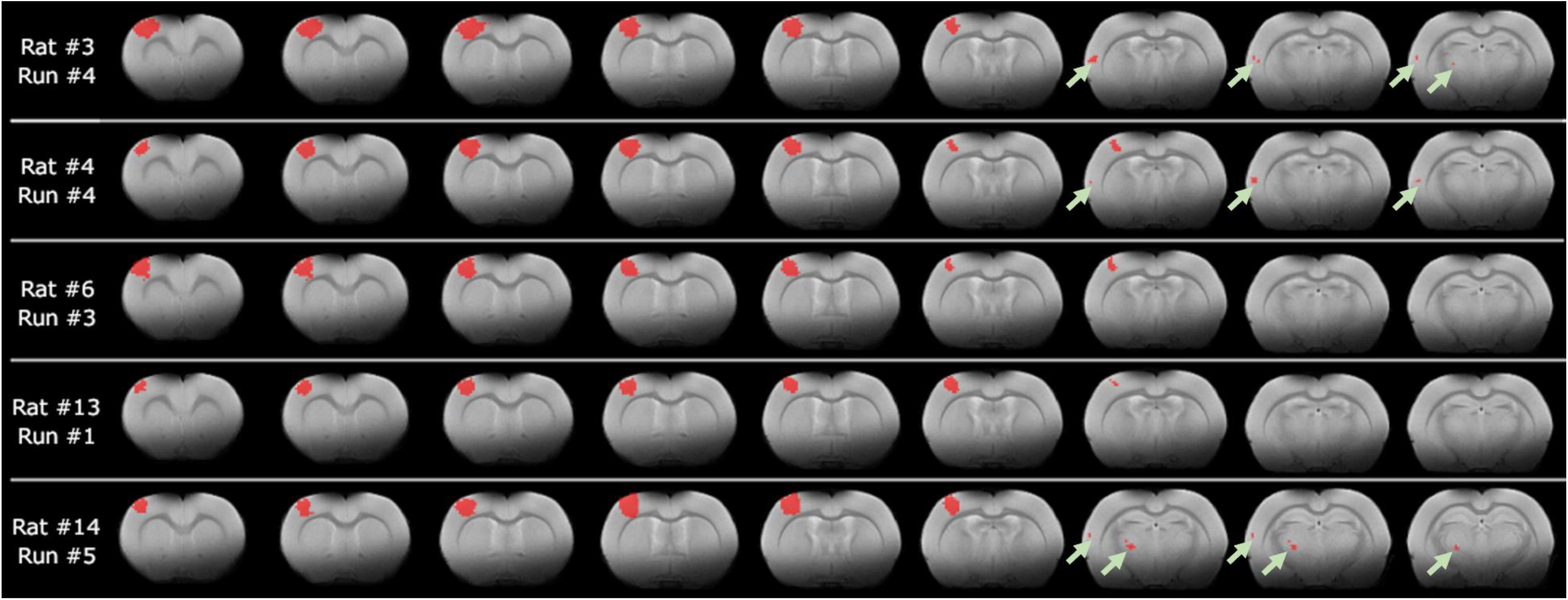
Examples of the extent of statistically significant voxels detected from the GLM analysis (p < 0.05, family-wise error correction) overlayed in red over T2w images for different rats and functional runs. The examples were chosen so as to illustrate the consistency of the positive BOLD response in the contralateral primary somatosensory cortex associated with unilateral forepaw stimulation, and the diversity of significant responses elsewhere in the brain, with sparser significant voxels distributions located in the secondary somatosensory cortex or the thalamus (green arrows).

**Figure S3:**
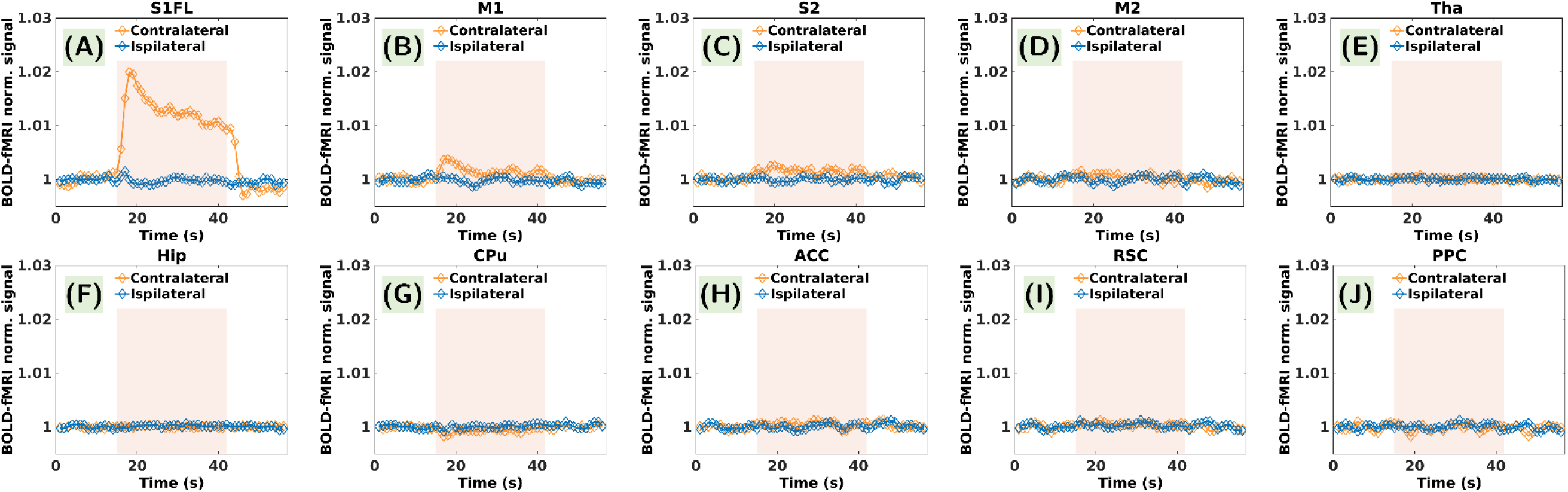
Contralateral and ipsilateral ROI-averaged BOLD response functions calculated by averaging across all epochs, rats and runs. The transparent orange overlay indicates the 28 s stimulation window. In contralateral S1FL a substantially fast initial progression characteristic of rodents can be observed. Within the 28-second stimulation window, a rapid surge followed by an overshoot reaching a peak of 2.0% above the baseline can be noticed. Subsequently, a gradual drop to 1% above the baseline unfolded over the next 25 s and was sustained up to stimulus termination. The signal returns to baseline after 3 s, and a noticeable poststimulus undershoot concludes the dynamic sequence of signal fluctuations. The contralateral M1 and S2 response functions display a noticeable signal increase during stimulation, while contralateral CPu presents a small decrease.

**Figure S4:**
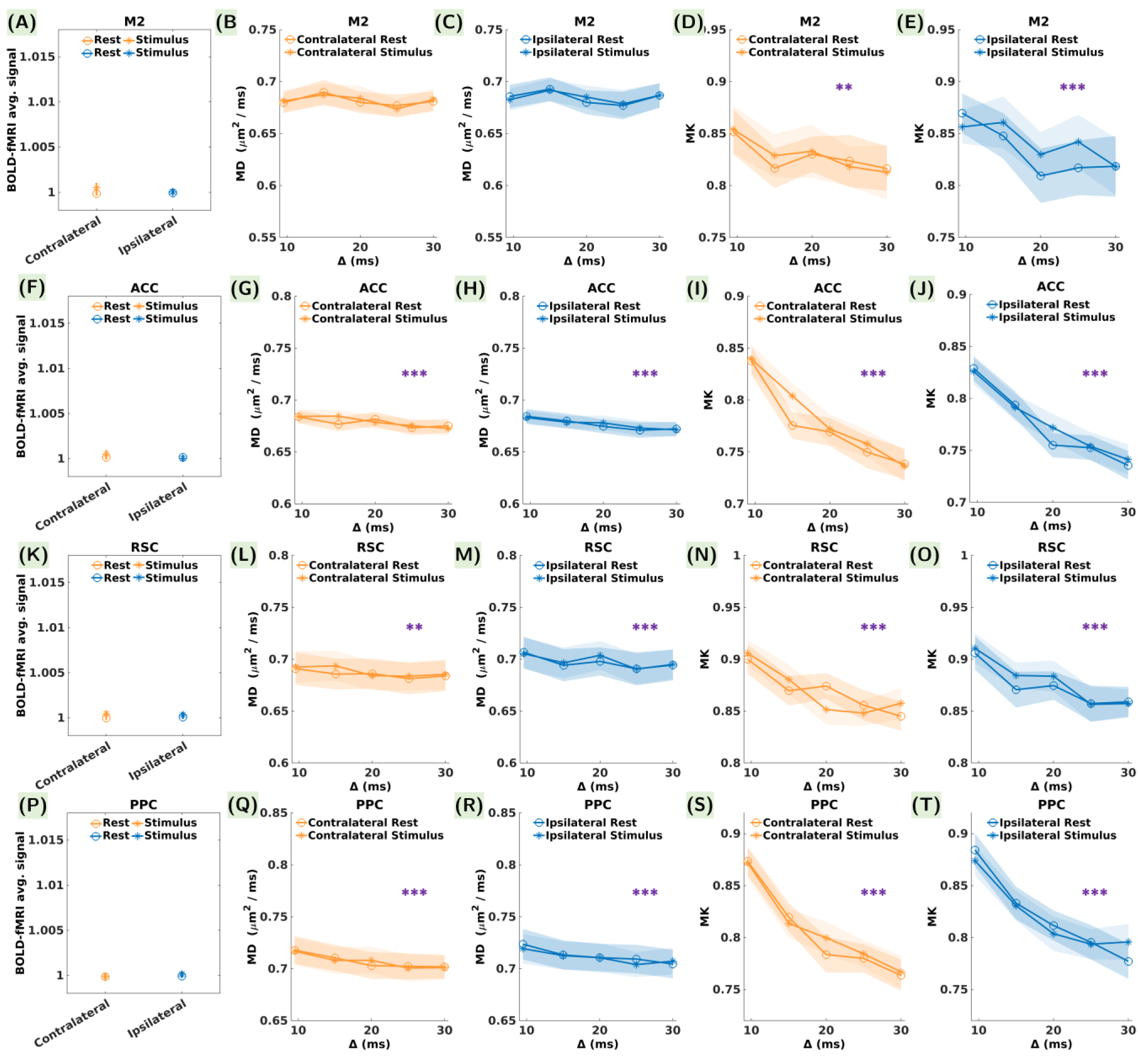
Average BOLD signal, MD and MK during the rest and stimulus conditions in control cortical brain regions (A-E) M2, (F-J) ACC, (K-O) RSC, and (P-T) PPC, contralaterally (blue) and ipsilaterally (orange). The error bars represent the standard error calculated over 24 measurements in each condition for BOLD and over 24 measurements in each condition and each diffusion time for MD and MK. No statistically significant differences were noticed between the rest and stimulus conditions for BOLD, MD or MK. Purple asterisks in the MD and MK plots indicate statistically significant differences between the diffusion time values. P-values are reported as: * p < 0.05, ** p < 0.01, *** p < 0.001 (FDR correction for multiple comparisons).

**Table S1:**
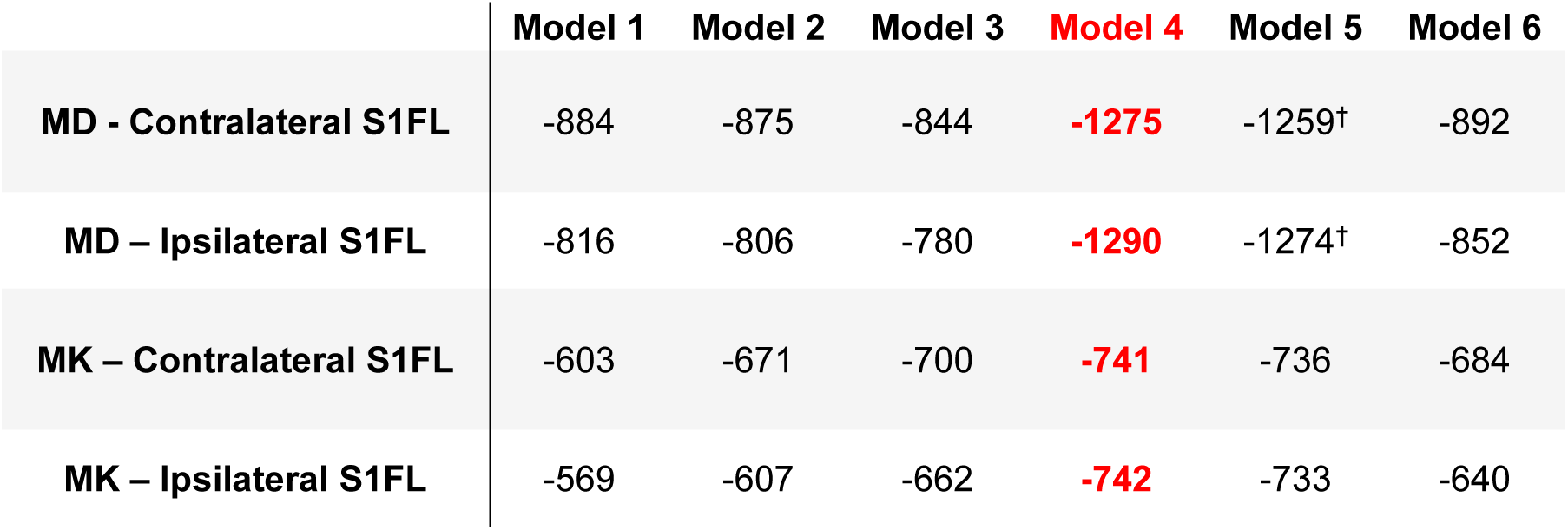
Mixed models tested on the dMRI data. Six different models with increasing complexity were tested on the MD and MK quantifications in S1FL. The model with the lowest Bayesian Information Criterion (BIC), in this case, Model 4 was selected and applied on all the other ROIs. We opted to gradually increase the complexity of the models. Thus, our initial model (Model 1) involved regressing the response variable against a constant term. By excluding predictor variables from this first model, we established a generic baseline against which we could evaluate the performance of all subsequent models. The next model (Model 2) included only our two main regressors (rest vs. stimulus and the diffusion time) as fixed effects, with no random effects. We then introduced the run index as a random effect in addition to the two main regressors (Model 3). Models 4 and 5 incorporated both the run and the subject indexes as random effects, once by nesting the subject index within the run number (Model 4), and once by nesting the run index within the subject index (Model 5). Finally, all four variables were considered as fixed effects in Model 6. The ^†^ symbol denotes a singular fit.

**Table S2:**
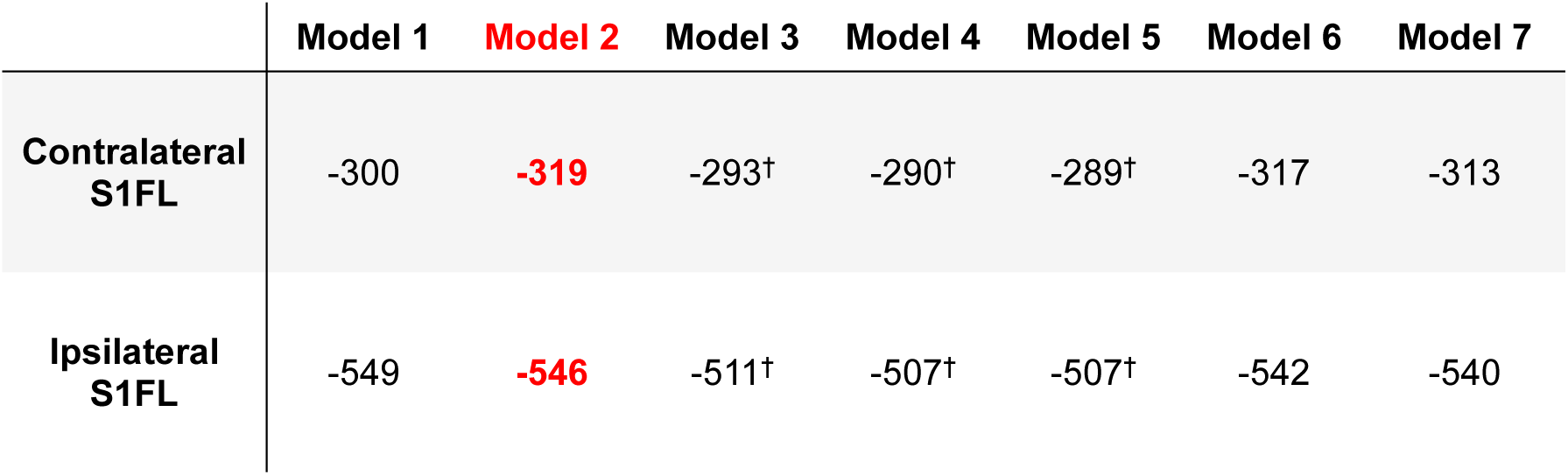
Mixed models tested on the BOLD signal. Seven different models with increasing complexity were tested on the MD and MK quantifications in S1FL. The model with the lowest Bayesian Information Criterion (BIC) was selected, in this case Model 2, and applied on all the other ROIs. We opted to gradually increase the complexity of the models. Thus, our initial model (Model 1) involved regressing the response variable against a constant term. By excluding predictor variables from this first model, we established a generic baseline against which we could evaluate the performance of all subsequent models. The next model (Model 2) included our main regressor (rest vs. stimulation) as a fixed effect, with no random effects. We then introduced the run index as a random effect in addition to the two main regressors (Model 3). Models 4 and 5 incorporated both the run and the subject indexes as random effects, once by nesting the subject index within the run number (Model 4), and once by nesting the run index within the subject index (Model 5). Next, the run index was considered as a fixed effect along with the main regressor (Model 6), and finally both the run and the subject indexes were considered as fixed effects (Model 7). The ^†^ symbol denotes a singular fit.

**Table S3:**
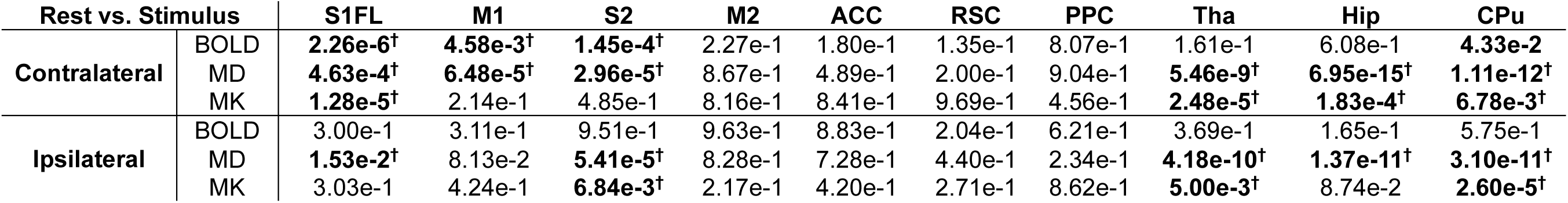
Results of the statistical analyses for rest vs. stimulus differences. Uncorrected p-values for statistically significant differences between the rest and stimulus conditions for BOLD, MD and MK in all the investigated ROIs contralaterally and ipsilaterally. In bold the significant p-values (< 0.05) are highlighted, and the dagger (**^†^**) symbol indicates statistically significant differences that survived FDR correction for multiple comparisons.

**Table S4:**
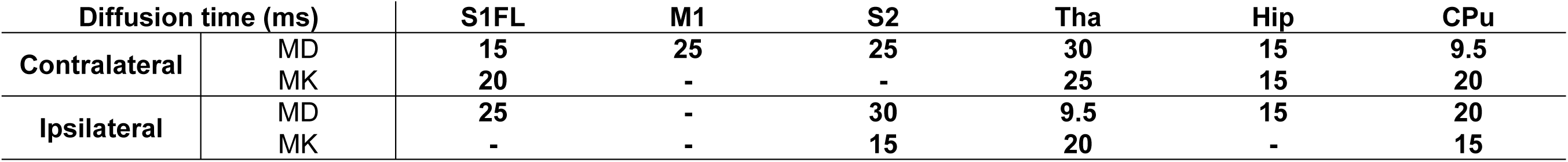
Diffusion time values corresponding to maximum MD and MK change during stimulation. Diffusion time values correspond to the maximum absolute changes in MD and MK amplitudes during stimulation reported in Table 1. Most of the investigated brain regions displayed a maximum change in MD and MK at the longer diffusion times probed in this study.

**Table S5:**
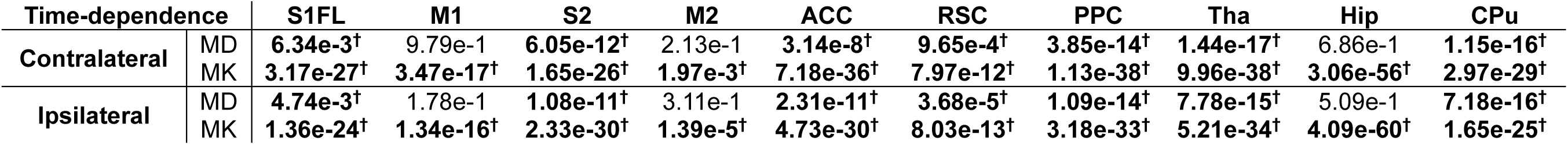
Results of the statistical analyses for differences between the various diffusion time values. Uncorrected p-values for statistically significant differences between the diffusion time values for MD and MK in all the investigated ROIs contralaterally and ipsilaterally. In bold the significant p-values (< 0.05) are highlighted, and the dagger (**^†^**) symbol indicates statistically significant differences that survived FDR correction for multiple comparisons.

