## Supplementary Material for "Somatosensory-evoked response induces extensive diffusivity and kurtosis changes associated with neural activity in rodents"

*Andreea Hertanu

Department of Radiology, Lausanne University Hospital (CHUV)

Rue du Bugnon 46, 1011 Lausanne

Phone : + 41 21 314 6020

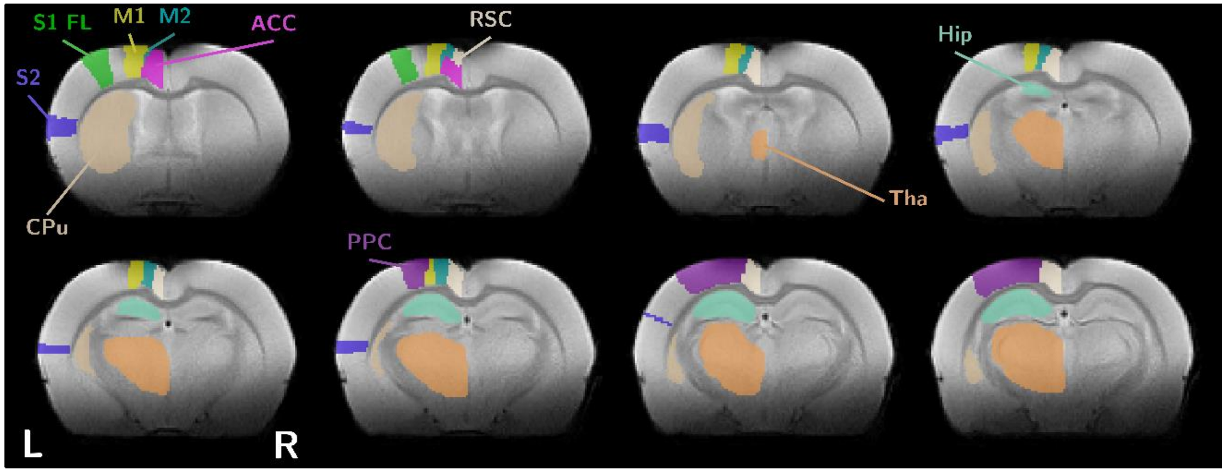


**Figure S1:** Illustration of the segmentation of various areas investigated in this study on slices spanning from the rostral the caudal brain: primary somatosensory cortex, forelimb area (S1FL), secondary somatosensory cortex (S2), primary motor cortex (M1), secondary motor cortex (M2), cingulate cortex (ACC), retrosplenial cortex (RSC), posterior parietal cortex (PPC), thalamus (Tha), striatum (CPu) and hippocampal subfields (Hip).

**Table S1: Mixed models tested on the dMRI data** - Six different models with increasing complexity were tested on the MD and MK quantifications in S1FL. The model with the lowest Bayesian Information Criterion (BIC), in this case, Model 4 was selected and applied on all the other ROIs. We opted to gradually increase the complexity of the models. Thus, our initial model (Model 1) involved regressing the response variable against a constant term. By excluding predictor variables from this first model, we established a generic baseline against which we could evaluate the performance of all subsequent models. The next model (Model 2) included only our two main regressors (rest vs. stimulus and the diffusion time) as fixed effects, with no random effects. We then introduced the run index as a random effect in addition to the two main regressors (Model 3). Models 4 and 5 incorporated both the run and the subject indexes as random effects, once by nesting the subject index within the run number (Model 4), and once by nesting the run index within the subject index (Model 5). Finally, all four variables were considered as fixed effects in Model 6. The † symbol denotes a singular fit.

| | Model 1 | Model 2 | Model 3 | **Model 4** | Model 5 | Model 6 |
|---|---|---|---|---|---|---|
| **MD - Contralateral S1FL** | -884 | -875 | -844 | **-1275** | -1259† | -892 |
| **MD – Ipsilateral S1FL** | -816 | -806 | -780 | **-1290** | -1274† | -852 |
| **MK – Contralateral S1FL** | -603 | -671 | -700 | **-741** | -736 | -684 |
| **MK – Ipsilateral S1FL** | -569 | -607 | -662 | **-742** | -733 | -640 |

**Table S2: Mixed models tested on the BOLD signal** - Seven different models with increasing complexity were tested on the MD and MK quantifications in S1FL. The model with the lowest Bayesian Information Criterion (BIC) was selected, in this case Model 2, and applied on all the other ROIs. We opted to gradually increase the complexity of the models. Thus, our initial model (Model 1) involved regressing the response variable against a constant term. By excluding predictor variables from this first model, we established a generic baseline against which we could evaluate the performance of all subsequent models. The next model (Model 2) included our main regressor (rest vs. stimulation) as a fixed effect, with no random effects. We then introduced the run index as a random effect in addition to the two main regressors (Model 3). Models 4 and 5 incorporated both the run and the subject indexes as random effects, once by nesting the subject index within the run number (Model 4), and once by nesting the run index within the subject index (Model 5). Next, the run index was considered as a fixed effect along with the main regressor (Model 6), and finally both the run and the subject indexes were considered as fixed effects (Model 7). The † symbol denotes a singular fit.

| | Model 1 | Model 2 | Model 3 | Model 4 | Model 5 | Model 6 | Model 7 |
|---|---|---|---|---|---|---|---|
| **Contralateral S1FL** | -300 | **-319** | -293† | -290† | -289† | -317 | -313 |
| **Ipsilateral S1FL** | -549 | **-546** | -511† | -507† | -507† | -542 | -540 |

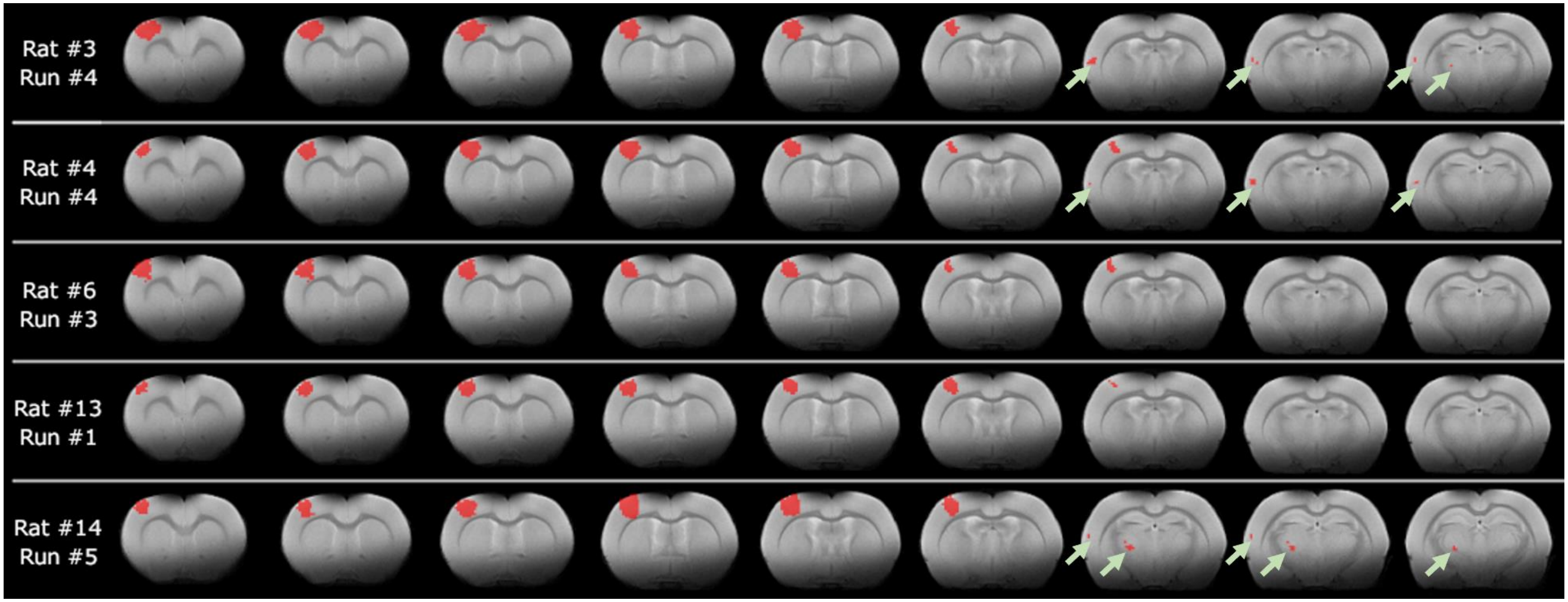


**Figure S2:** Examples of the extent of statistically significant voxels detected from the GLM analysis ($p < 0.05$, family-wise error correction) overlayed in red over $T_2$w images for different rats and functional runs. The examples were chosen so as to illustrate the consistency of the positive BOLD response in the contralateral primary somatosensory cortex associated with unilateral forepaw stimulation, and the diversity of significant responses elsewhere in the brain, with sparser significant voxels distributions located in the secondary somatosensory cortex or the thalamus (green arrows).

**Table S3**: **Results of the statistical analyses for rest vs. stimulus differences** - Uncorrected p-values for statistically significant differences between the rest and stimulus conditions for BOLD, MD and MK in all the investigated ROIs contralaterally and ipsilaterally. In bold the significant p-values (< 0.05) are highlighted, and the dagger (†) symbol indicates statistically significant differences that survived FDR correction for multiple comparisons.

| Rest vs. Stimulus | | S1FL | M1 | S2 | M2 | ACC | RSC | PPC | Tha | Hip | CPu |
|---|---|---|---|---|---|---|---|---|---|---|---|
| **Contralateral** | BOLD | **2.26e-6†** | **4.58e-3†** | **1.45e-4†** | 2.27e-1 | 1.80e-1 | 1.35e-1 | 8.07e-1 | 1.61e-1 | 6.08e-1 | **4.33e-2** |
| | MD | **4.63e-4†** | **6.48e-5†** | **2.96e-5†** | 8.67e-1 | 4.89e-1 | 2.00e-1 | 9.04e-1 | **5.46e-9†** | **6.95e-15†** | **1.11e-12†** |
| | MK | **1.28e-5†** | 2.14e-1 | 4.85e-1 | 8.16e-1 | 8.41e-1 | 9.69e-1 | 4.56e-1 | **2.48e-5†** | **1.83e-4†** | **6.78e-3†** |
| **Ipsilateral** | BOLD | 3.00e-1 | 3.11e-1 | 9.51e-1 | 9.63e-1 | 8.83e-1 | 2.04e-1 | 6.21e-1 | 3.69e-1 | 1.65e-1 | 5.75e-1 |
| | MD | **1.53e-2†** | 8.13e-2 | **5.41e-5†** | 8.28e-1 | 7.28e-1 | 4.40e-1 | 2.34e-1 | **4.18e-10†** | **1.37e-11†** | **3.10e-11†** |
| | MK | 3.03e-1 | 4.24e-1 | **6.84e-3†** | 2.17e-1 | 4.20e-1 | 2.71e-1 | 8.62e-1 | **5.00e-3†** | 8.74e-2 | **2.60e-5†** |

**Table S4**: **Diffusion time values corresponding to maximum MD and MK change during stimulation** – Diffusion time values correspond to the maximum absolute changes in MD and MK amplitudes during stimulation reported in Table 1. Most of the investigated brain regions displayed a maximum change in MD and MK at the longer diffusion times probed in this study.

| **Diffusion time (ms)** | | **S1FL** | **M1** | **S2** | **Tha** | **Hip** | **CPu** |
|---|---|---|---|---|---|---|---|
| **Contralateral** | MD | **15** | **25** | **25** | **30** | **15** | **9.5** |
| | MK | **20** | **-** | **-** | **25** | **15** | **20** |
| **Ipsilateral** | MD | **25** | **-** | **30** | **9.5** | **15** | **20** |
| | MK | **-** | **-** | **15** | **20** | **-** | **15** |

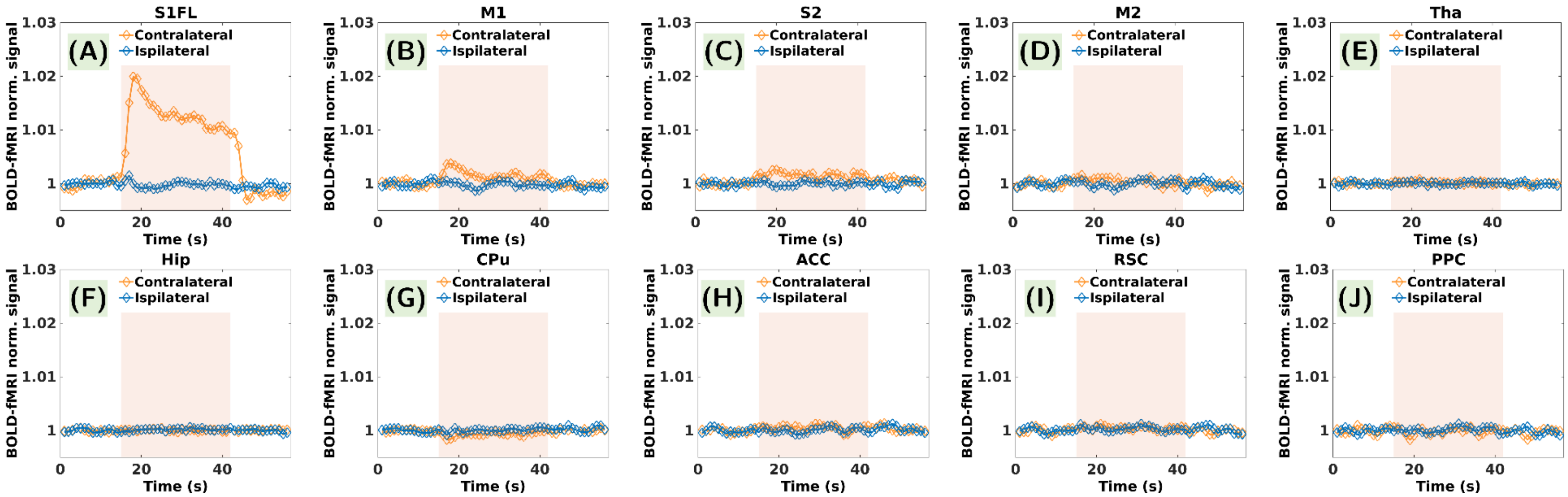


**Figure S3:** Contralateral and ipsilateral ROI-averaged BOLD response functions calculated by averaging across all epochs, rats and runs. The transparent orange overlay indicates the 28 s stimulation window. In contralateral S1FL a substantially fast initial progression characteristic of rodents can be observed. Within the 28-second stimulation window, a rapid surge followed by an overshoot reaching a peak of 2.0% above the baseline can be noticed. Subsequently, a gradual drop to 1% above the baseline unfolded over the next 25 s and was sustained up to stimulus termination. The signal returns to baseline after 3 s, and a noticeable poststimulus undershoot concludes the dynamic sequence of signal fluctuations. The contralateral M1 and S2 response functions display a noticeable signal increase during stimulation, while contralateral CPu presents a small decrease.

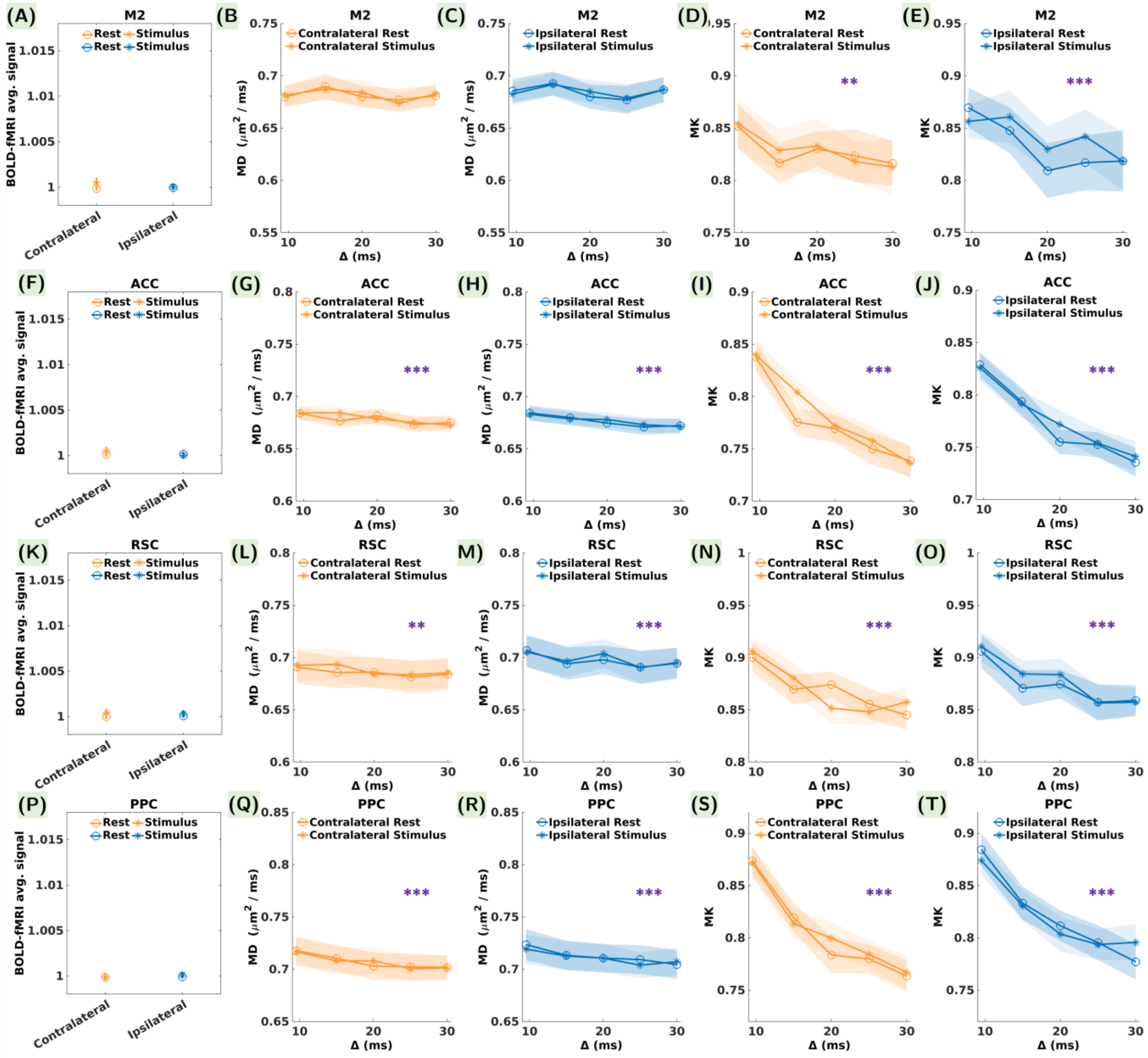

**Figure S4:** Average BOLD signal, MD and MK during the rest and stimulus conditions in control cortical brain regions (A-E) M2, (F-J) ACC, (K-O) RSC, and (P-T) PPC, contralaterally (blue) and ipsilaterally (orange). The error bars represent the standard error calculated over 24 measurements in each condition for BOLD and over 24 measurements in each condition and each diffusion time for MD and MK. No statistically significant differences were noticed between the rest and stimulus conditions for BOLD, MD or MK. Purple asterisks in the MD and MK plots indicate statistically significant differences between the diffusion time values. P-values are reported as: * $p < 0.05$, ** $p < 0.01$, *** $p < 0.001$ (FDR correction for multiple comparisons).

**Table S5**: **Results of the statistical analyses for differences between the various diffusion time values** - Uncorrected p-values for statistically significant differences between the diffusion time values for MD and MK in all the investigated ROIs contralaterally and ipsilaterally. In bold the significant p-values (< 0.05) are highlighted, and the dagger (†) symbol indicates statistically significant differences that survived FDR correction for multiple comparisons.

| **Time-dependence** | | **S1FL** | **M1** | **S2** | **M2** | **ACC** | **RSC** | **PPC** | **Tha** | **Hip** | **CPu** |
|---|---|---|---|---|---|---|---|---|---|---|---|
| **Contralateral** | MD | **6.34e-3†** | 9.79e-1 | **6.05e-12†** | 2.13e-1 | **3.14e-8†** | **9.65e-4†** | **3.85e-14†** | **1.44e-17†** | 6.86e-1 | **1.15e-16†** |
| | MK | **3.17e-27†** | **3.47e-17†** | **1.65e-26†** | **1.97e-3†** | **7.18e-36†** | **7.97e-12†** | **1.13e-38†** | **9.96e-38†** | **3.06e-56†** | **2.97e-29†** |
| **Ipsilateral** | MD | **4.74e-3†** | 1.78e-1 | **1.08e-11†** | 3.11e-1 | **2.31e-11†** | **3.68e-5†** | **1.09e-14†** | **7.78e-15†** | 5.09e-1 | **7.18e-16†** |
| | MK | **1.36e-24†** | **1.34e-16†** | **2.33e-30†** | **1.39e-5†** | **4.73e-30†** | **8.03e-13†** | **3.18e-33†** | **5.21e-34†** | **4.09e-60†** | **1.65e-25†** |
